# Sequential and dynamic coding of water-sucrose categorization in rat gustatory cortices

**DOI:** 10.1101/2024.04.19.590344

**Authors:** Germán Mendoza, Esmeralda Fonseca, Hugo Merchant, Ranier Gutierrez

## Abstract

The gustatory system enables our conscious perception of sweetness, allowing us to distinguish sweet solutions from water. However, the neural mechanisms underlying this categorization process in rats remain poorly understood. We addressed this question by designing a novel sucrose categorization task in which rats classified varying sucrose concentrations against water. We found that in the anterior insular cortex (aIC) and orbitofrontal cortex (OFC), neural activity primarily encoded the categorical distinction between sucrose and water, rather than specific sucrose concentrations. Notably, aIC neurons encoded this distinction faster than OFC neurons. Conversely, the OFC slightly preceded the aIC in encoding choice information, although both cortices maintained parallel encoding of the rat’s choices. The encoding of sensory and categorical decisions was dynamic and sequential, forming a sequence of encoding neurons throughout the trial. These findings reveal that sucrose categorization relies on dynamic coding sequences in the neuronal activity of the aIC and OFC rather than static, long-lasting (sustained) neural representations. This dynamic coding, supported by single-cell population decoding and principal component analyses, suggests that the brain continuously updates its representation of sucrose categorization as new information emerges. Additionally, both the aIC and OFC rapidly encoded reward outcomes. Our data supports the notion that gustatory cortices employ sequential and dynamic coding to compute sensorimotor transformations, from taste detection to categorical taste decisions and reward processing.

**Graphical Abstract:** 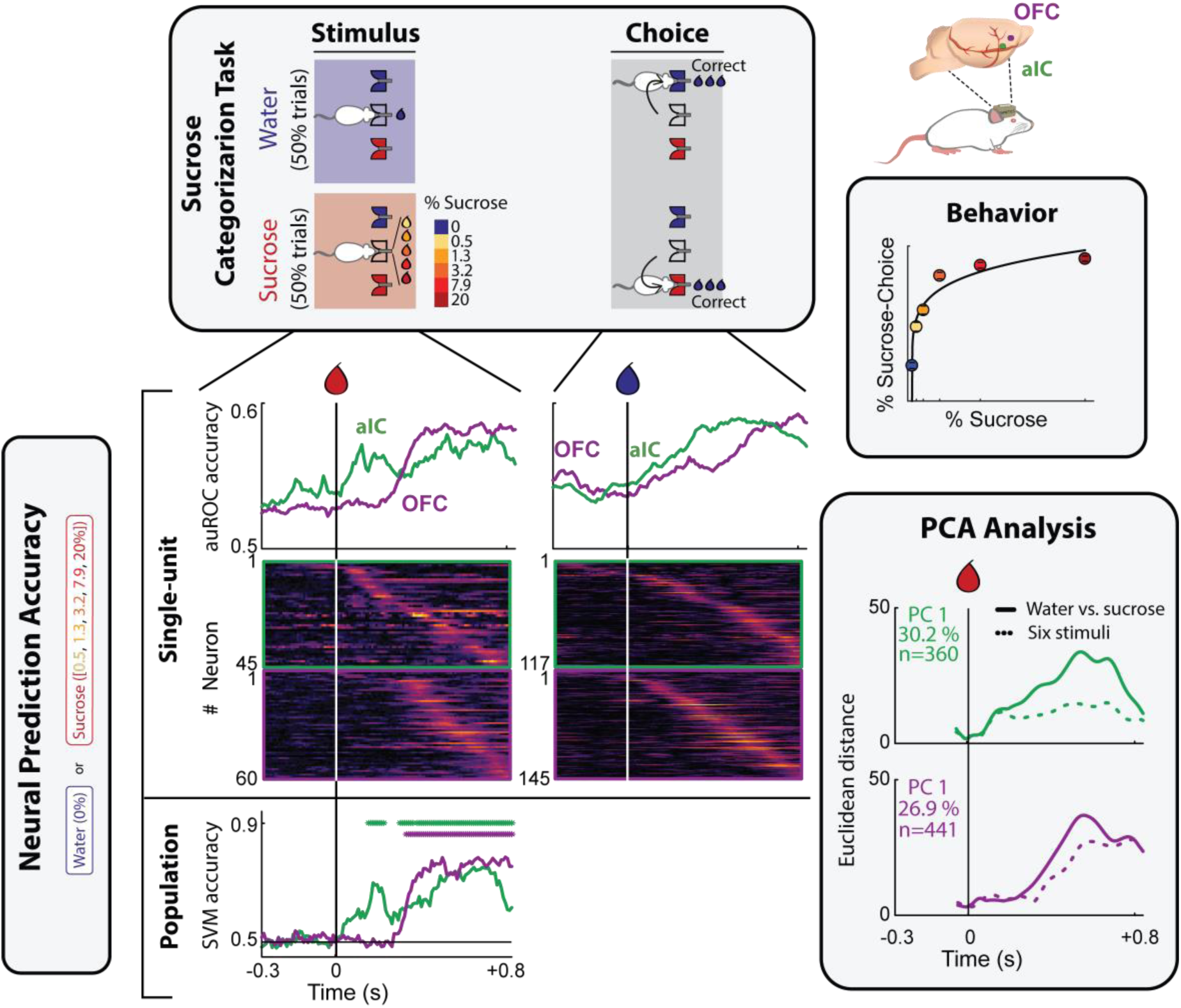

## Introduction

Sucrose, commonly found in table sugar and fruits, is a high-calorie disaccharide that most animals find palatable and elicits a conscious experience of sweetness in humans and likely in rodents. Its consumption can lead to obesity and elevate the risk of various obesity-related chronic diseases, including type 2 diabetes, heart disease, and stroke.^1^ Due to its medical relevance, extensive research has been conducted to understand its sensory and perceptual processing.^2–4^ The anterior Insular (aIC) and Orbitofrontal Cortices (OFC) function as primary and secondary taste cortices and evaluate food taste and value,^5,6^ including sweetness.^7,8^ Although most experiments studying the neural mechanism of sweetness perception have focused on passive taste processing,^6,9–12^ some studies have investigated how the brain actively uses gustatory percepts to make decisions.^7,13–15^ For instance, we previously discovered that aIC and OFC neurons in rats discriminating between different sucrose concentrations sparsely encode sweetness intensity, choice behavior, and reward outcomes.^7^ Nevertheless, the neural mechanisms of sweetness perception that drive the voluntary consumption of sucrose are not yet fully understood. For example, whether the brain can precisely extract and encode the different sucrose concentrations in food remains unclear. Prior studies have reported some degree of monotonic encoding of sweet tastant concentrations in the gustatory cortices.^6,7,9–12^ Still, only a few studies^7,8^ have quantified the significance of these parametric neural representations for behavior, and none have contrasted them with a hypothesis of a more categorical, or less precise, encoding in active decision-making.

Conversely, several rodent and primate studies have observed dynamic population activity to encode behaviorally relevant variables.^16–18^ In these dynamic codes, neurons with similar tuning properties convey relevant information at different and relatively brief time windows within a trial, generating a sequence or cascade of neural activity.^17,19^ Such sequential activity can retain or encode information for extended periods, effectively serving as a working memory substrate.^19,20^ Dynamic codes are also characterized by changes in the tuning properties of some neurons, with neurons encoding different information in different epochs of the trial.^17,19^ Population-level analyses based on dimension reduction techniques, such as PCA, have been used to study these neural dynamics across various species and tasks.^21–23^ Interestingly, no studies have reported the existence of dynamic population encoding in the rat’s aIC and OFC during sweet tastant processing, nor how these encoding patterns are reflected in the low-dimensional representations of population activity.

In the present study, we used single-neuron recordings, neural population decoding, and dimensionality reduction techniques to compare encoding dynamics in the aIC and OFC during a sucrose categorization task. In this task, rats were required to sample a drop of water-based liquid and then report whether the stimulus was plain water or contained sucrose. Sucrose could be presented in one of five concentrations, ranging from 0.5% to 20%. Our findings revealed that the distinction between the identity/quality of the tastant (water vs. sucrose), rather than sucrose intensity (i.e., specific concentrations), was the most robust stimulus-related information encoded in the population activity of aIC and OFC neurons. We also observed that neurons in both cortices dynamically encoded stimulus-related information and the rat’s choices, with significant information carried by different neurons at specific time points, resulting in reproducible encoding sequences. Some neurons exhibited a dynamic shift in their encoding patterns, transitioning between perceptual decisions, outcomes, or task variables. Finally, we found that all the main features observed in the activity of single neurons or with population decoding techniques were also reflected in a linear low-dimensional state space of neuronal population activity. This flexibility in cortical coding highlights the adaptability of the gustatory system in actively processing and integrating complex taste information.

## Behavioral Results

### Rats could solve the sucrose categorization task

Eleven male Sprague-Dawley rats were trained in a novel sucrose categorization task. The task involved classifying various sucrose solutions as “Water” or “Sucrose” based on the concentration of sucrose. The trial structure was as follows: rats started each trial by licking a central spout (2-3 times) to receive a 10 µL water or sucrose solution (stimulus delivery epoch). Next, rats chose a lateral spout (water or sucrose-associated spouts) and emitted one dry lick (Choice epoch; **Figure 1A**). A correct choice triggered the delivery of three water drops as a reward, one per subsequent lick; an incorrect choice resulted in no reward. During each trial, one of six stimuli was delivered: water (0% sucrose) or sucrose (0.5%, 1.3%, 3.2%, 7.9%, and 20%). Water was used as the stimulus in 50% of the trials per experimental session, while the remaining sucrose concentrations were evenly distributed in the other 50%.

**Figure 1.**
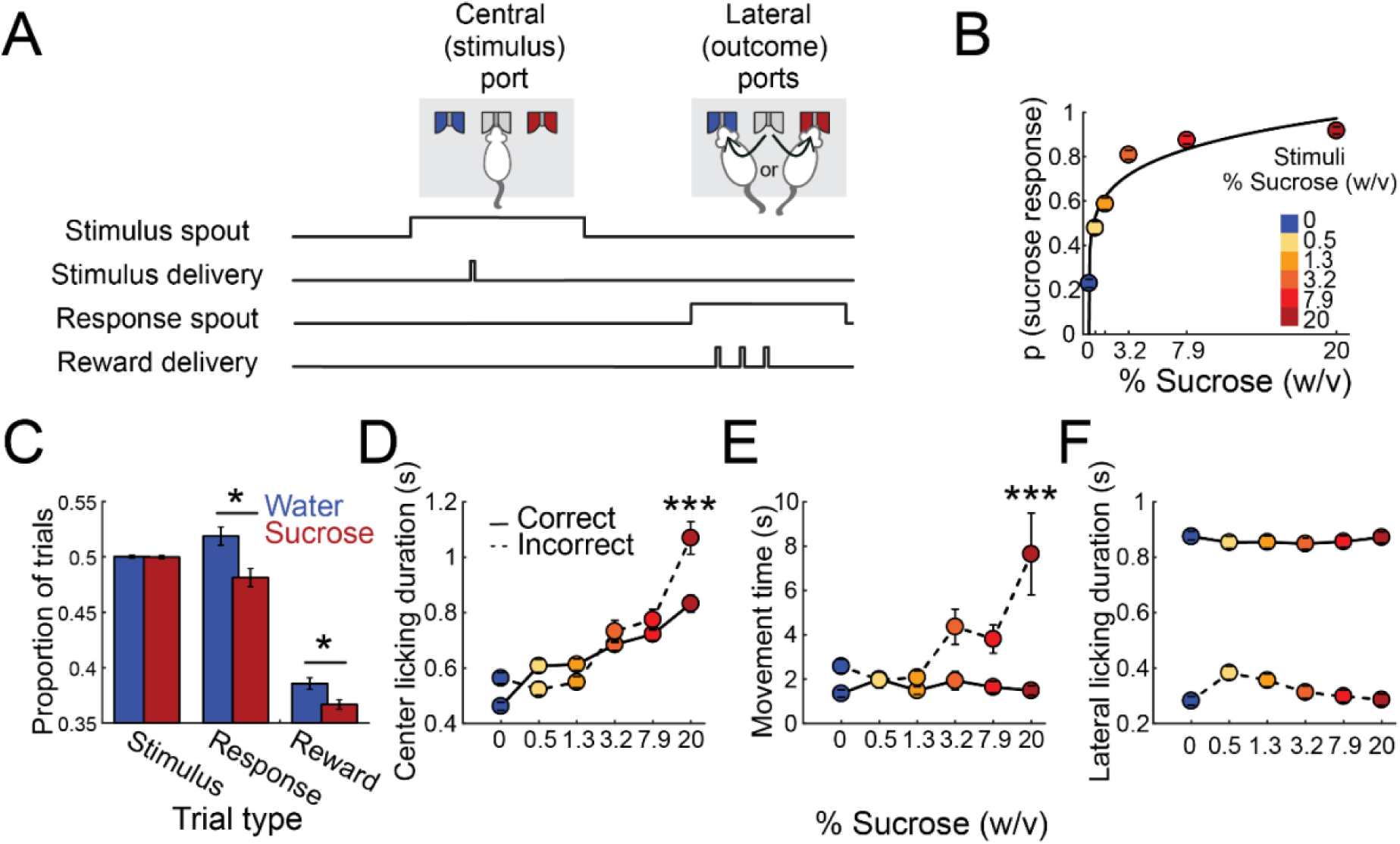
Water-sucrose categorization task in rats. (**A**) Schematic diagram of the behavioral setup. The rat is presented with three licking spouts: a central spout (stimulus port) and two lateral spouts (outcome ports). At the beginning of each trial, the rat licks the central spout to receive a 10 µL drop of liquid as a stimulus. The stimulus is one of six different sucrose concentrations dissolved in water: 0 (water), 0.5, 1.3, 3.2, 7.9, and 20% w/v. The rat must then make a water/sucrose choice by licking the water- or sucrose-associated lateral spout. If the correct lateral spout is chosen, the rat receives three drops of water as a reward after one empty lick. The session is composed of 50% water trials. The other sucrose concentrations were equally distributed over the remaining trials. (**B**) Probability of selecting the sucrose-associated lateral spout (sucrose response) for each stimulus. The black line depicts the power fitting (R^2^ = 0.87). Each stimulus is represented according to the color code displayed on the right sidebar. (**C**) Trial distribution. The proportion of trials with water (blue) or sucrose (red) is shown in the left panel. The proportion of trials in which the rat chose the water (blue) or sucrose spout (red) is shown in the middle panel. The proportion of correct (rewarded) water (blue) or sucrose (red) responses is depicted in the right panel. Asterisks indicate significant differences according to one-tailed t-tests. (**D**) Time spent licking the central spout, from stimulus delivery until the last lick, before responding to a lateral spout. The color code in panel (**B**) indicates the different stimuli delivered. Continuous and dashed lines indicate correct and incorrect trials, respectively. Asterisks indicate significant differences across correct and incorrect trials within sucrose concentrations according to a two-way ANOVA and a Tukey-Kramer pairwise comparison test (see Table 2 for all the pairwise comparisons). (**E**) Latency to the lateral spout’s first lick after the central spout’s last lick. Same conventions as in (**D**). See Table 3 for all the pairwise comparisons. (**F**) Licking duration at the lateral spout. Data from each subject’s sessions were included and presented as mean ± SEM. Same conventions as in (D). See Table 4 for pairwise comparisons.

We plotted the average psychometric performance to demonstrate that rats could categorize gustatory stimuli based on sucrose concentration (**Figure 1B**). The graph depicts the probability of rats perceiving and reporting each stimulus as sucrose. As expected, the probability of a sucrose choice increased steadily with higher sucrose concentrations. From the curve fitting to the average psychometric performance, we determined that the absolute threshold, defined as the predicted concentration at which rats would respond to the sucrose choice spout in 50% of the trials, was 0.35%. From the actual tested concentrations, 0.5% and 1.3% were the nearest to the predicted absolute threshold. Their probability of responding to the sucrose spout was closer to 50%, indicating that these concentrations were the hardest to detect from the actual stimuli set. Thus, rats could not reliably distinguish these two concentrations from water. The presentation of water and sucrose stimulus trials was balanced; however, rats made a significantly higher proportion of ‘water’ choices compared to ‘sucrose’ choices (0.518 ± 0.081 vs. 0.481 ± 0.081, respectively; t-test, t_(100)_ = 2.29, p = 0.012; **Figure 1C**). The proportion of correct/rewarded trials was also higher for water-response than sucrose-response trials (0.385 ± 0.052 vs. 0.366 ± 0.043; t-test, t_(100)_ = 2.18, p = 0.015, **Figure 1C**). Altogether, these results suggest that the increased proportion of water-response trials was due to the difficulty in correctly categorizing the lowest sucrose concentrations (0.5% and 1.3%).

### Sucrose palatability was reflected as increased dry licking intervals to the stimulus spout

Once the tastant is delivered in gustatory perceptual decision-making tasks, the time spent between the first lick to consume the stimulus and the last dry lick to the central spout is a measurement that reflects the palatability or hedonic value of the gustatory cue.^24^ In our categorization task, stimulus spout-licking duration increased as a function of sucrose concentration; this response pattern was maintained during correct trials and incorrect trials (Two-way ANOVA with sucrose concentration and trial’s outcome as factors and licking duration in the center spout as the dependent variable. F_concentration(5)_ = 93.855, p < 0.001; F_outcome(1)_ = 13.084, p < 0.001; F_interaction(5)_ = 12.524, p < 0.001. See **Supplementary Table 1** for all the pairwise comparisons; **Figure 1D**). Thus, despite the trial outcome, a preserved palatability effect was exhibited throughout the stimulus spout licking. That is, as palatability increases, so does licking duration. Overall, these results suggest that, in taste-guided decisions, the duration of dry licks emitted after stimulus sampling might reflect the subject’s palatability and perception.

### Movement time during correctly categorized trials did not contain palatability information

Interestingly, rats exhibited similar movement times (time between the last stimulus-spout lick and the first response-spout lick) across all stimuli during correct responses (Two-way ANOVA with sucrose concentration and trial’s outcome as factors and movement time as the dependent variable. F_concentration(5)_ = 7.57, p < 0.001; F_outcome(1)_ = 46.22, p < 0.001; F_interaction(5)_ = 8.15, p < 0.001. See **Supplementary Table 2** for all the pairwise comparisons; **Figure 1E**), which is consistent with other psychophysical taste tasks.^24^ Thus, the movement was ballistic and confident in correct trials and did not reflect taste palatability. However, movement times significantly increased during error trials for the highest sucrose concentration (See **Supplementary Table 2** for all the pairwise comparisons; **Figure 1E**). This suggests that rats hesitated longer during incorrect choices, particularly when licking high-sucrose solutions. It is important to note that when rats moved from the central to the lateral spout, they kept updating their choices; thus, this epoch was rich in decision-making information.

### Rats rapidly detect reward delivery from omission

For correct trials, rats exhibited sustained licking following reward delivery (three drops of water as a reward). They spent significantly more time licking the response spout after rewarded trials compared to non-rewarded ones (**Figure 1F**. Two-way ANOVA with sucrose concentration and trial’s outcome as factors and lateral licking duration as the dependent variable. F_concentration(5)_ = 3.11, p < 0.001; F_outcome(1)_ = 4407.42, p < 0.001; F_interaction(5)_ = 5.98, p < 0.001. See **Supplementary Table 3** for all the pairwise comparisons). These findings suggest that rats not only distinguished a reward but also rapidly stopped licking when no reward was delivered (error trial). This allowed them to rapidly restart another trial, complete more trials, and thus collect more rewards throughout the session. The latter confirms that rats can swiftly adapt their behavior based on reward outcomes.

## Electrophysiological results

### A single drop of tastant triggers a ramp-up of cell recruitment

We recorded 400 neurons from the aIC and 449 neurons from the OFC in the left hemisphere of rats performing the sucrose categorization task (**Figure S1**). To investigate the single-neuron temporal dynamics of task-information encoding on a trial-by-trial basis, we applied a moving window Signal Detection Theory-based analysis to the firing rates of each neuron (100 ms time windows^7,15,25^ with moving steps of 10 ms^26^). We found similar proportions of neurons encoding the stimulus, the choice, and the outcome in the aIC and OFC (**Figure S2**). In both gustatory cortices, the stimulus variable was represented by the smallest subset, with 11.6% and 14.22% of stimulus-encoding neurons in the aIC and OFC, respectively (z = - 1.109, p = 0.267), followed by the choice-encoding subset with 30.15% and 34.36% of cells (z = -1.278, p = 0.200), and finally, the outcome-encoding subset with 54.12% and 54.74% of neurons in the aIC and OFC, respectively (z = -1.175, p = 0.857). Thus, a single (quantal) drop of taste stimulus triggered an avalanche of activity that gradually recruited an increasing number of cortical neurons, from stimulus delivery to the reward outcome, encompassing all relevant task variables that may give rise to trial structure representation.^7,27^ Furthermore, we found relatively small neuronal subpopulations that encoded more than one variable. In the following paragraphs, we describe these results in detail.

### Neurons in aIC and OFC encoded the categorical distinction between water and sucrose

To identify neurons that categorically encoded stimulus information, we assessed differences in neural activity between water-stimulus (sucrose 0%) and sucrose-stimulus (sucrose 0.5, 1.3, 3.2, 7.9, 20%) trials per single neuron recorded. Only correct trials were included in the analysis. Firing rate distributions for water- and sucrose-stimulus trials were calculated per time bin for each cell. The distribution overlap was measured by computing the area under the Receiver Operating Characteristic (auROC) curve, referred to as stimulus auROC. The auROC curve values range between 0 and 1; values closer to the edges, 0 or 1, indicate fully segregated distributions, while a 0.5 value indicates complete overlap. In this case, the auROC reflects the proportion of water or sucrose trials decoded from single-neuron activity on a trial-by-trial basis. It is important to note that, in categorization tasks, a trial-by-trial correlation between the stimulus and the subject’s choice during correct trials could emerge, making it challenging to dissociate stimulus-related neural activity from the choice. We addressed this problem by computing an additional auROC curve that does not consider the received stimulus (water or sucrose) but what the subject perceived according to their choice/response; this curve will be referred to as the “choice probability index.” We considered stimulus encoding when statistically significant stimulus auROC time-bins were also higher than the corresponding choice probability index. Neurons with five or more consecutive time bins meeting these criteria were classified as stimulus-encoding cells. The window containing the largest significant auROC value was considered the best categorical stimulus-encoding time window (see **Figure 2A**, cyan rectangles).

**Figure 2.**
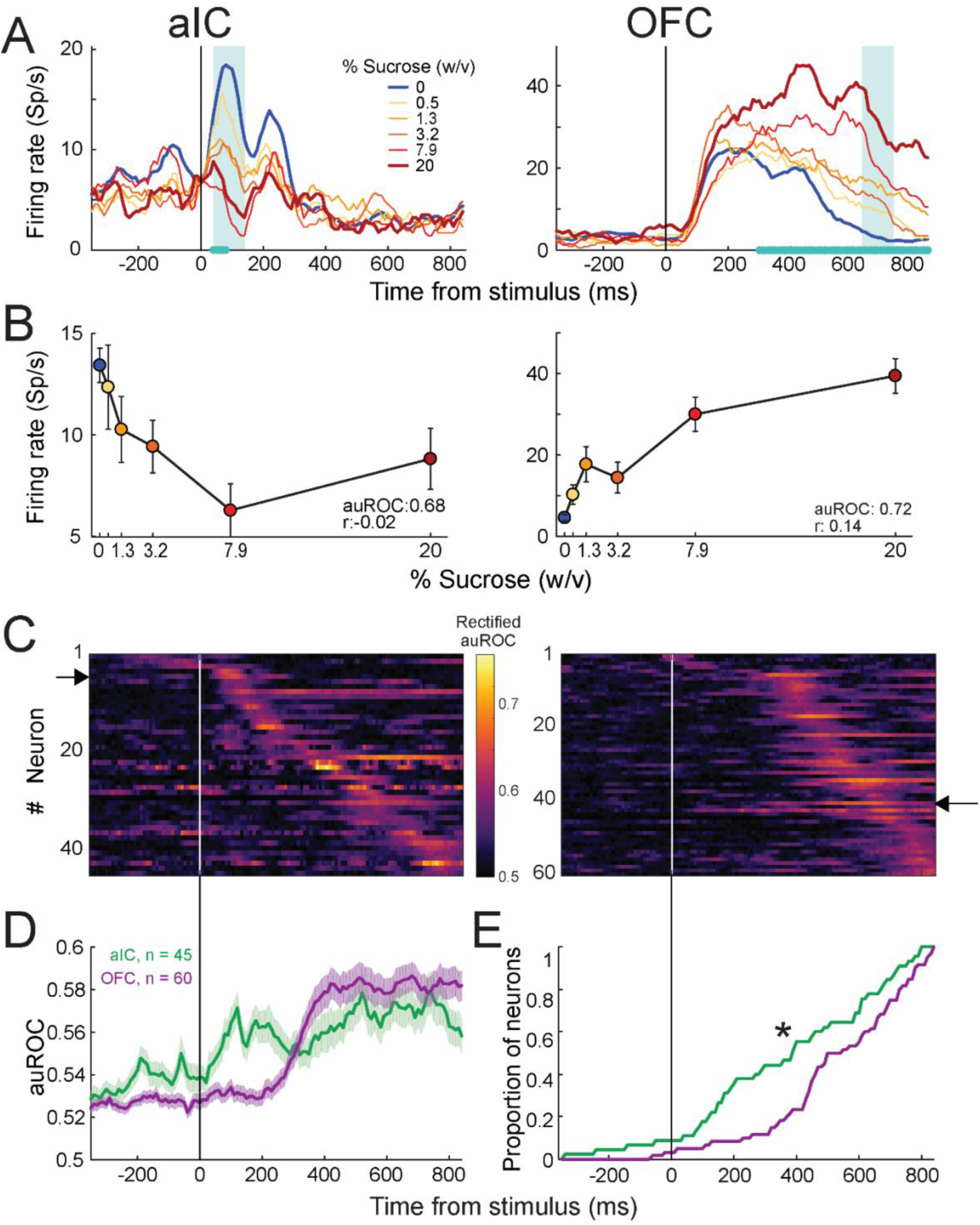
aIC and OFC neurons encode the presence or absence of sucrose. (**A**) Representative examples of aIC (left) and OFC (right) neurons that respond differentially to the absence (water) and presence (scale of reds) of sucrose. The activity is aligned (time = 0 s) with sucrose delivery in the central spout. The color code shows sucrose concentration in the upper right corner. The cyan asterisks on the abscissa indicate the time windows with significant encoding according to a permutation test (10,000 permutations; see methods). The cyan rectangle marks the time window with the maximum significant area under the auROC value. Figure S2 shows the proportion of recruited neurons per task epoch. (**B**) Averaged neuronal activity during the best stimulus-encoding time window (cyan rectangle in (**A**) as a function of sucrose concentration. (**C**) Subpopulations of aIC (left) and OFC (right panel) neurons that encode sucrose detection. The color map shows the auROC values (color code on the right) for each significant stimulus-encoding neuron (y-axis) across time aligned to stimulus delivery (x-axis). Neurons are ordered according to the time of maximum auROC value. Black arrows indicate the neurons shown in (**A**). The stimulus was encoded in sequence for both regions, but it was slightly more sequential in aIC (SqI: 0.749; PE: 0.981; TS: 0.571) than OFC (SqI: 0.695; PE: 0.939; TS: 0.515). (**D**) Mean auROC values per time bin for the aIC (green) and OFC (purple) subpopulations shown in (**C**). (**E**) Cumulative proportion to time of the auROC peak values of the populations shown in (**C**). The black asterisk indicates a significant difference between distributions as determined by a two-sample Kolmogorov-Smirnov test, * p < 0.01. Data are presented as mean ± SEM.

We found a small group of categorical stimulus-encoding cells (water vs. sucrose) in both the aIC (45 out of 388, 11.6%) and OFC (60 out of 422, 14.22%). The percentage of stimulus-encoding neurons in each cortex was not significantly different (z = -1.109, p = 0.267; **Figure S2**), consistent with previous findings suggesting that taste coding is sparse and distributed.^7,28^ **Figure 2A** shows examples of auROC-identified aIC and OFC stimulus-encoding neurons. The aIC neuron showed phasic categorical stimulus-selective activity around 50 ms after tastant delivery; it rapidly responded to the water stimulus and exhibited a transient inhibition after sucrose stimulus delivery. In contrast, the OFC neuron rapidly ramped up (around 50 ms) but exhibited stimulus-selective activity after 200 ms from stimulus delivery, with a higher firing rate during sucrose-stimulus trials compared to water-stimulus trials. Time intervals containing the most significant neural activity difference between water- and sucrose-stimulus trials, which is the best time window for categorical stimulus encoding (see cyan asterisks over the time axis, **Figure 2A**), occurred at distinct moments after stimulus delivery. Both cells also significantly encoded the intensity (all concentrations) of the stimulus with different temporal signatures, as shown by the cyan-shaded rectangle indicating the best time window for intensity stimulus encoding (**Figure 2A**). The mean activity elicited by each stimulus concentration during this best time window is shown in **Figure 2B**. Interestingly, all stimulus-encoding cells encoded both types of information in different non-overlapping time windows. Altogether, our results demonstrate that both gustatory cortices represented stimulus information in a temporal, dynamic, distributed, and sparse manner.

### Transient sequences of stimulus-encoding neurons in the aIC and OFC reliably represent the categorical distinction between sucrose and water

We wondered whether the encoding period of each stimulus-related cell would sequentially encompass the entire stimulus epoch. To visualize the sequential encoding of each cell belonging to the stimulus-encoding subset, we plotted the color-coded auROC values across the stimulus epoch from -250 ms to +800 ms aligned to stimulus delivery. Neurons were sorted based on their auROC peak latency. Note that the information encoding pattern in both areas is dynamic (i.e., each neuron exhibited higher auROC values during short time windows), and the encoding windows of different cells are scattered across time (**Figure 2C**). Accordingly, the stimulus-encoding peak time did not occur simultaneously across cells but was distributed after stimulus delivery. We computed an adapted version of the Sequentiality Index (SqI) developed by^16^ to determine whether aIC and OFC stimulus-encoding cells represent stimulus information through neural activity sequences homogeneously distributed across neurons and span the stimulus epoch in time. The original index is calculated with neural population firing rates and is bounded between 0 and 1. The SqI approaches 1 when the peak times of each neuron homogeneously tile the entire duration of interest, and one neuron is active at every moment in time, with the temporal fields being non-overlapping. The index is computed from Peak Entropy (PE), which measures the entropy of the distribution of peak times across the entire neural population, and Temporal Sparsity (TS), which measures the entropy of the distribution of normalized activity in any bin. Here, we applied the Sql to the auROC values shown in **Figure 2C** (see Methods). This analysis examines the sequential nature of information encoding in the aIC and OFC. In this way, PE reflects how homogeneously the encoding periods of the different neurons tile the entire analyzed period, with values close to one for more homogenous tiling and zero otherwise. TS measures how much of the activity at each point in time can be attributed to a single neuron, with values of zero and one corresponding to overlapped and non-overlapped encoding periods, respectively. We found that the stimulus was sequentially encoded in both cortical regions, but the sequentiality was slightly greater in the aIC (SqI: 0.749; PE: 0.981; TS: 0.571) than in the OFC (SqI: 0.695; PE: 0.939; TS: 0.515). Also, the relatively high SqI depended mainly on the PE, not the TS, indicating that the encoding periods homogeneously covered the analyzed stimulus epoch but with a medium overlapping of the encoding periods of different cortical neurons.

We compared them in different trial subsets to test whether the neural encoding sequences are reproducible and not a random product. First, auROC values were obtained from two randomly partitioned sets of trials. This resulted in two matrices of auROC values per cortical area. Next, we sorted the cells in matrix one by their latencies to peak auROC for each cortical area, used the same indexes to sort the second matrix, and computed the Mean-Squared Error (MSE) between both sorted matrices to measure pattern similarity. For aIC and OFC neurons, the MSE between splits was 0.0036 and 0.0021, respectively. A permutation test showed that these relatively low errors in the aIC and OFC populations could not be obtained by chance (p < 0.001), supporting the notion that stimulus-encoding sequences are highly reproducible.

Another main observation in the color-coded maps of **Figure 2C** is that the encoding of stimulus information by OFC neurons is delayed by about 200 ms compared to that of aIC neurons. This can be seen in **Figure 2D**, which depicts the average auROC across statistically significant neurons for both cortical areas. The mean auROC of the aIC population tends to increase rapidly (> 50 ms) and steadily after stimulus delivery. In contrast, stimulus selectivity for the OFC remained low until 300 ms, when it abruptly increased, outperforming the aIC. **Figure 2E** shows the cumulative distributions of the proportion of neurons that reached their peak auROC values across time bins. Rapid neuronal recruitment in the aIC followed by the OFC can be seen. A two-sample Kolmogorov-Smirnov test on the time bins at which aIC and OFC neurons reached their peak auROC values (p < 0.01) supports this observation. These results revealed that stimulus-encoding aIC cells were recruited first, followed by the OFC stimulus-encoding subset.

Furthermore, aIC and OFC stimulus-encoding cells represented their information in the shape of stimulus-selective neural sequences. These patterns rose from the most substantial stimulus information (greatest auROC values for water stimulus vs. sucrose stimulus) contained in distinct, short time windows across cells. Of note, aIC stimulus-selective sequences started earlier and were more robust than the OFC ones.

### At the population level, stimulus-encoding cells in the aIC and OFC encoded the categorical distinction between water and sucrose more robustly than between specific concentrations

At the single-neuron level, aIC and OFC stimulus-encoding cells represented information in a categorical (water vs sucrose) manner (**Figure 2**). auROC analyses were performed to determine the categorical stimulus-encoding subset. Next, using a Support Vector Machine (SVM) decoder, we evaluated whether the already auROC-identified stimulus-encoding cells also contained stimulus information at the neuronal population level. We also assessed the impact of using linear vs. non-linear kernels on SVM stimulus-decoding performance. For each cortical area, we fed the activity of the auROC-identified Stimulus-encoding neurons, one vs. one, into an 8-fold cross-validated SVM classifier that decoded the identity of the stimulus (the class) presented in each trial. The algorithm splits the multi-class classification into a single binary classification problem per each pair of classes. The proportion of in-fold trials correctly classified by the model trained in the out-of-fold trials was considered a measure of decoding performance. We used linear, Gaussian, or third-order polynomial kernels to compare SVM performances. The first kernel is preferred for classification problems with linearly separable data; Gaussian and polynomial kernels perform non-linear data transformations and are employed when a non-linear relationship between features exists.^29^ The linear kernel was not outperformed by the Gaussian or polynomial kernels, suggesting that the data were linearly separable (**Figure S3**); thus, we will focus on the linear-kernel SVM results. Stimulus-decoding accuracy was generally higher for the OFC than for the aIC and remained below 0.4 (the chance level for six stimuli is 0.166; **Figure S3A**). Also, the difference in encoding latencies was measured, with the aIC leading the OFC. For comparison, we trained the SVM to classify the trials as water stimulus vs. sucrose stimulus using the activity of the same selective neurons (**Figure S3B**). The decoding accuracy in both cortical regions reached around 0.75 (the chance level for two stimuli is 0.5; **Figure S3B**). Thus, the SVM performed better in decoding the water-sucrose distinction than in decoding the six stimuli in the aIC and OFC. Altogether, aIC and OFC stimulus-encoding cells represented categorical and intensity information at the single unit and population levels, with the categorical feature containing the most information.

One caveat to this analysis is that it focused on the activity of a small number of cells that had already been screened for their single-unit selectivity to the water vs. sucrose stimulus conditions. Therefore, we additionally used ANOVA on all recorded single units to identify the neurons with significant firing rate differences between the six stimulus concentrations and ran the same SVM decoding analysis. This approach allowed us to extract all possible information from the neural populations. We excluded neurons with significant choice encoding—according to the choice probability index from the analysis—to minimize potential contamination from neural signals associated with the subject’s choices (see Methods). A total of 154 aIC and 181 OFC neurons were identified. These expanded neural populations resulted in similar qualitative results (**Figures 3A and 3C**), suggesting that the neurons identified by the auROC analyses have as much information as the extended populations. More importantly, these observations corroborate our findings that the categorical distinction between stimuli (water vs. sucrose) was more robust than the gradual intensity representation of all stimulus concentrations.

**Figure 3.**
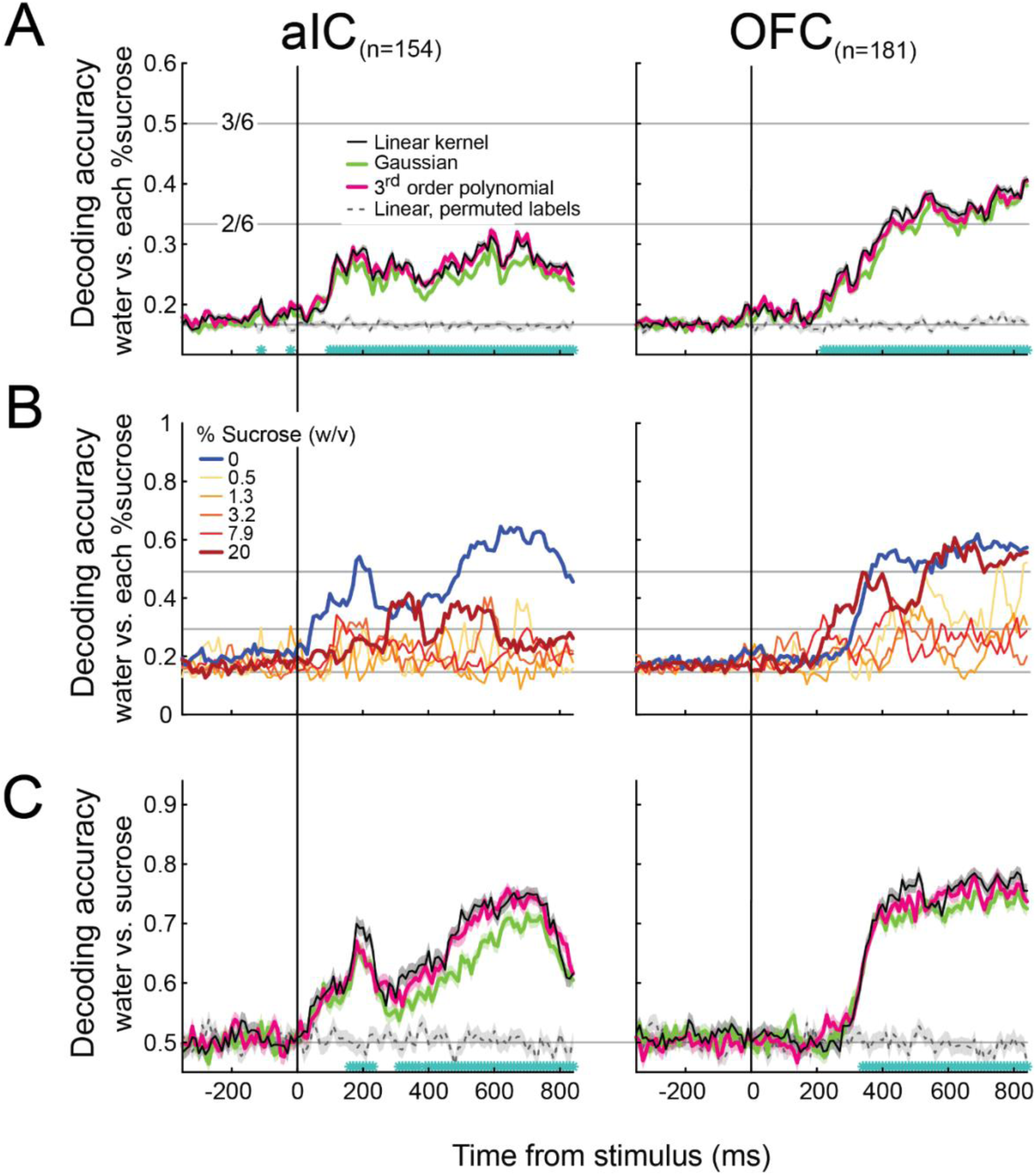
The qualitative distinction between water and sucrose, rather than the gradual change in intensity of the stimulus, is the most robust sensory-related information in the firing rate of the neural populations in aIC and OFC. (**A**) SVM accuracy to decode the six stimuli across time depending upon the type of kernel used. Decoders were trained with the activity of the extended population of neurons detected with ANOVA, which contains significant firing rate differences between the six stimulus concentrations and is aligned to stimulus delivery. The black, green, and pink lines depict the linear, Gaussian, and third-order polynomial kernels. The gray dashed line indicates the performance of the linear kernel after the stimulus labels have been randomly permuted. The horizontal light-gray continuous lines indicate the decoding performance for correctly identifying 1 out of 6 stimuli (chance), 2 out of 6 stimuli, and 3 out of 6 stimuli, from bottom to top. Cyan asterisks indicate bins where significant differences between the linear SVM and the linear permuted SVM were found (one-tailed t-test, p<= 0.0001, Bonferroni corrected). Data are presented as mean ± SEM. (**B**) The proportion of correct classifications for each stimulus by the linear-kernel SVM is shown in (**A**). (**C**) Decoding accuracy of SVMs using a linear, Gaussian, or third-order polynomial kernel to decode the water or sucrose stimulus from the activity of aIC (left) and OFC (right) neurons. The light-gray horizontal line indicates chance performance (0.5). Same conventions as in (**A**). See **Figure S3**, which shows the same analysis on only the significant auROC neurons.

To determine the influence of each stimulus over the linear SVM decoder accuracy (**Figure 3A**, black line), we computed the proportion of trials in which each concentration was correctly classified (**Figure 3B**). A decoder accuracy value of 0.5 indicates that the decoder correctly classified that concentration in half of the trials. It can now be seen that water-stimulus trials had the highest proportion of correct classifications for the aIC and OFC. In the OFC, though, the highest sucrose concentration also exhibited a comparable proportion of correctly classified trials. Overall, in these gustatory cortices, both water and 20% sucrose were the stimuli most likely to be classified correctly. In sum, in a taste-based perceptual decision-making task, intensity information of the easiest-to-detect stimuli is represented more prominently at the aIC and OFC population levels. By using not only auROC-identified stimulus-encoding cells but also a larger subset of neurons, we extended our results and confirmed that identity (i.e., water vs sucrose)—not intensity—was the most critical stimulus information contained in the aIC and OFC populations.

To further investigate whether the performance of the linear-kernel decoder trained with aIC or OFC neural activity aligns more closely with gross discrimination (water vs. sucrose) than with fine discrimination (differentiating sucrose concentrations), confusion matrices were computed (**Figure S4**). We calculated the mean-squared error (MSE) between the matrices derived from aIC (**Figure S4C**) or OFC (**Figure S4D**) neural activity and the theoretical matrices for fine (**Figure S4A**) and gross (**Figure S4B**) discrimination patterns. The results revealed that both aIC and OFC decoding patterns exhibited a stronger resemblance to the gross categorization pattern than to the fine discrimination pattern (MSE aIC vs. fine: 0.11; MSE aIC vs. gross: 0.03; MSE OFC vs. fine: 0.08; MSE OFC vs. gross: 0.04).

### During the choice epoch, categorical decisions were sequentially represented by aIC and OFC neurons

Having found aIC and OFC subsets that contained information about stimulus identity at the single-unit and population levels during the stimulus epoch, we sought to determine whether the subject’s response/choice was represented during the response epoch (-300 ms to +800 ms after the last lick to the stimulus spout). We performed a sliding window auROC analysis to identify choice-encoding cells and compared the firing rates of water-response vs. sucrose-response trials. In total, 117 of 388 (30.15%) aIC cells and 145 of 422 (34.36%) OFC cells significantly encoded the perceptual judgment of the rats, regardless of the delivered stimulus (**Figure S2**). **Figure 4A** depicts representative choice-encoding neurons from the aIC and OFC. The aIC choice-encoding neuron exhibited a first peak of activity similar to all stimuli shortly after the rat exited the central port and most likely had already made a choice. This neuron also showed a second activity peak at around 400 to 600 ms, which was categorically choice-selective (see cyan asterisks; **Figure 4A**, left panel). This aIC neuron remained silent during water-response trials (see solid lines) but was active during sucrose-response trials (dashed lines), regardless of the stimulus delivered at the central port, including water stimulus (i.e., an incorrect trial; see blue dashed line). The OFC choice-encoding cell displayed a similar activity pattern (**Figure 4A**, right panel). **Figure 4B** depicts the neurometric curves (red) computed with the neural activity indicated by the cyan-shaded rectangles in **A** and the corresponding psychometric curves (black) calculated with the rat’s responses/choices within the corresponding experimental sessions. The chosen time window contains the closest correspondence between the neurometric and the psychometric curves on a trial-by-trial basis, indicating that the activity of both neurons is highly related to the rats’ decisions. From the 117 aIC choice-encoding neurons, 51 (43.6%) showed higher activity for water responses, 51 (43.6%) for sucrose responses, and 15 (12.8%) showed a shift in their response patterns during the trial (**Figure S5A,** left panels). For the 145 OFC choice-encoding neurons, 67 (46.2%) exhibited preferred activity for water responses, 49 (33.8%) preferred sucrose responses, and 29 (20%) showed a shift in their response patterns during the response epoch (**Figure S5A**, right panels). This evidence suggests that the taste system might also use dynamic choice encoding. We computed the SqI, PE, and TS from the choice auROC values during the choice epoch to determine whether choice-encoding neurons sequentially monitor this decision variable across time. **Figure 4C** illustrates the choice auROCs values of the choice-encoding aIC and OFC cells sorted based on their peak latencies. The SqI, PE, and TS indexes were 0.603, 0.96, and 0.376 for aIC and 0.53, 0.989, and 0.284 for OFC. These results suggest that choice, like stimulus, is represented slightly more sequentially in the aIC than in the OFC. We observed that these sequences were highly reproducible in different trial splits. For aIC and OFC neurons, the MSE between splits was 0.0036 and 0.0021, respectively, which were highly unlikely to result by chance (p < 0.001). The temporal pattern of choice-encoding cell auROC values shows that choice information increases as the subject leaves the stimulus spout (**Figure 4D**). Nonetheless, unlike stimulus encoding during the stimulus epoch, we found no significant differences in the temporal recruitment of choice cells in the aIC compared to the OFC (**Figure 4E**; two-sample Kolmogorov-Smirnov test, p = 0.07). **Figure 4E** shows an earlier slight bump in the number of choice-encoding cells in the OFC compared to the aIC (-200 to 100 ms), which was then surpassed by the aIC (400-800 ms) after the last central spout lick. Our results suggest that both taste cortices sequentially represented the subjects’ perceptual judgment on a trial-by-trial basis with a marked overlap in recruitment.

**Figure 4.**
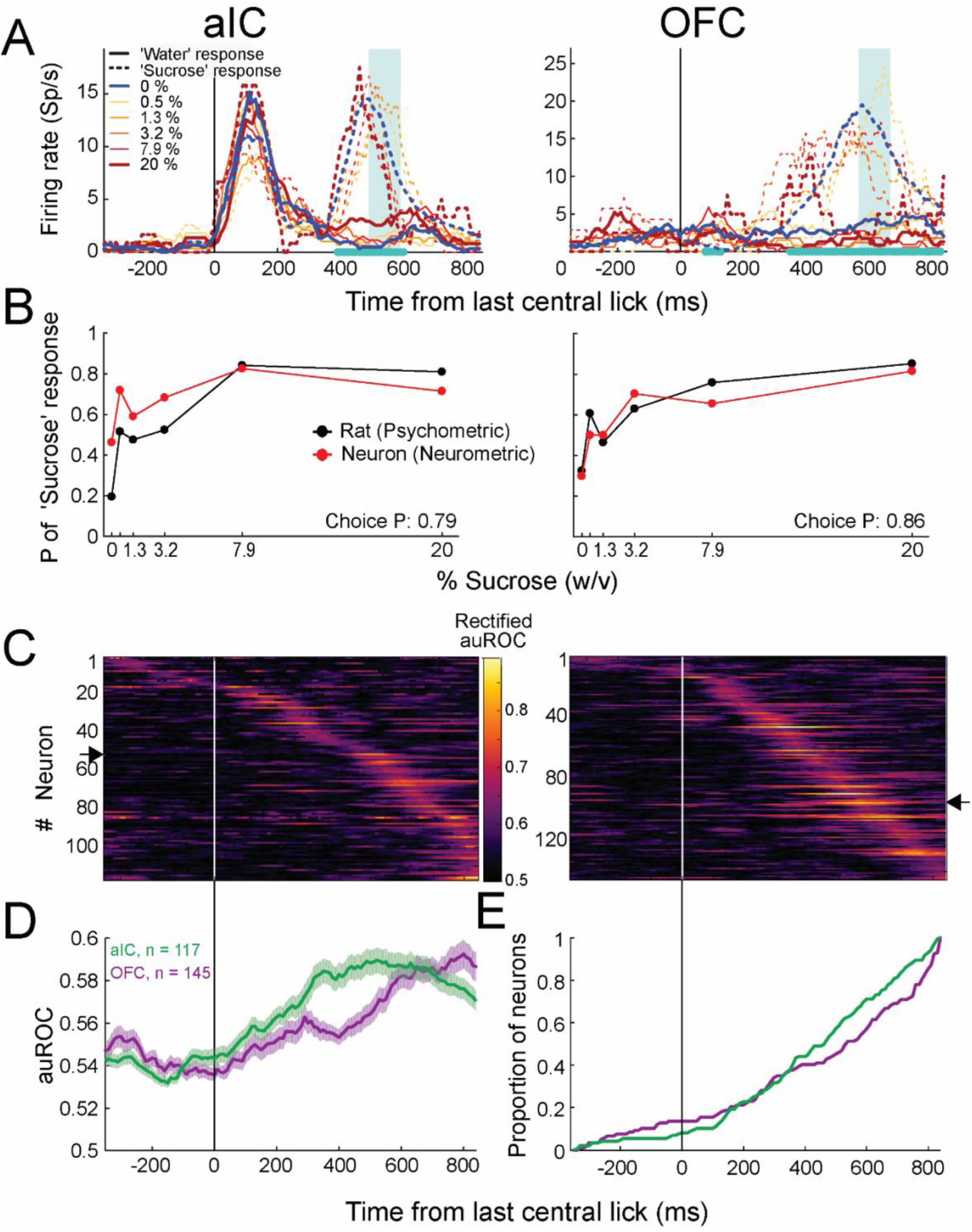
aIC and OFC neurons encode the categorical choice of the rats. **(A)** The mean firing rate of one aIC (left) and one OFC (right) neuron is selective to the animal’s choice. The activity is aligned (Time = 0 s) to the last lick in the central spout, encompassing the response epoch of the trial. Neural activity is separated by stimulus (color-coded) and choice: water or sucrose choice is marked by continuous and dashed lines, respectively. The cyan asterisks on the abscissa indicate the time windows with significant encoding according to a permutation test (10,000 permutations; see methods). The cyan rectangle marks the time window with the maximum significant area under the receiver operating characteristic curve [auROC] value. The aIC neuron exhibited a peak of activity around 400 to 600 ms only for the sucrose-response trials (dashed lines). The OFC neuron in the right panel also responded more to trials where the rat chose the sucrose response port. **(B)** Probabilities of ‘Sucrose’ response, obtained from the subject (psychometric; black) and predicted by the neuron (neurometric; red) as a function of the stimuli. The probabilities were calculated from the number of sucrose-response trials in the experimental session where neurons plotted in (**A**) were recorded. Decoded probabilities were calculated from the activity periods (cyan shaded) in (**A**) by classifying each trial as water or sucrose response according to an optimum firing rate threshold (see main text). **(C)** Rectified choice probability values per time bin of each neuron in the choice-encoding subpopulation of aIC (left) and OFC (right). Data aligned (Time = 0 s) to last central empty lick. (D) Average choice probability values of all choice-encoding neurons. **(E)** Cumulative proportion of choice-encoding cells that reached their peak auROC values across time bins. The same conventions as in **Figure 2**. Data presented as mean <u>±</u> SEM. See **Figure S5** for the average activity of neurons responding more to water and sucrose choices or responding to different choices at different time bins.

### Taste cortex neurons in both regions exhibited rapid and parallel encoding of reward outcomes

Finally, utilizing a similar auROC analysis, we quantified the neurons that distinguish reward delivery or omission during the outcome epoch, from -300 to +800 ms after the second lick to the choice spout (feedback time). We found outcome-encoding cells in both the aIC (210 of 388, 54.12%) and OFC (231 of 422, 54.74%) (**Figure S2**). These neurons exhibited selective activity for correct vs. incorrect trials, irrespective of the delivered stimulus. The outcome variable was represented by the largest subset in both GCs, compared to the stimulus- and choice-encoding subset sizes.

Next, we illustrate the activity of two representative outcome-encoding neurons (**Figure 5A**) whose activity was aligned with the outcome epoch start (feedback time). Both cells exhibited higher activity during correct trials than during error trials (solid vs. dotted lines). Remarkably, the activity pattern during all six stimulus trial types (0-20%) was highly similar within the same outcome condition (correct vs. incorrect). aIC cell activity (left panel) was also related to the rat’s responses before feedback. Specifically, the aIC exhibited higher activity for sucrose choices, regardless of whether they were correct or incorrect (see arrow in **Figure 5A**, left panel). After feedback, the activity became more influenced by the outcome: higher activity for correct trials and lower activity for incorrect trials (see the cyan-shaded rectangle). Similarly, the OFC outcome-encoding cell responded more when the reward was delivered than when it was omitted (**Figure 5A**, left panel).

**Figure 5.**
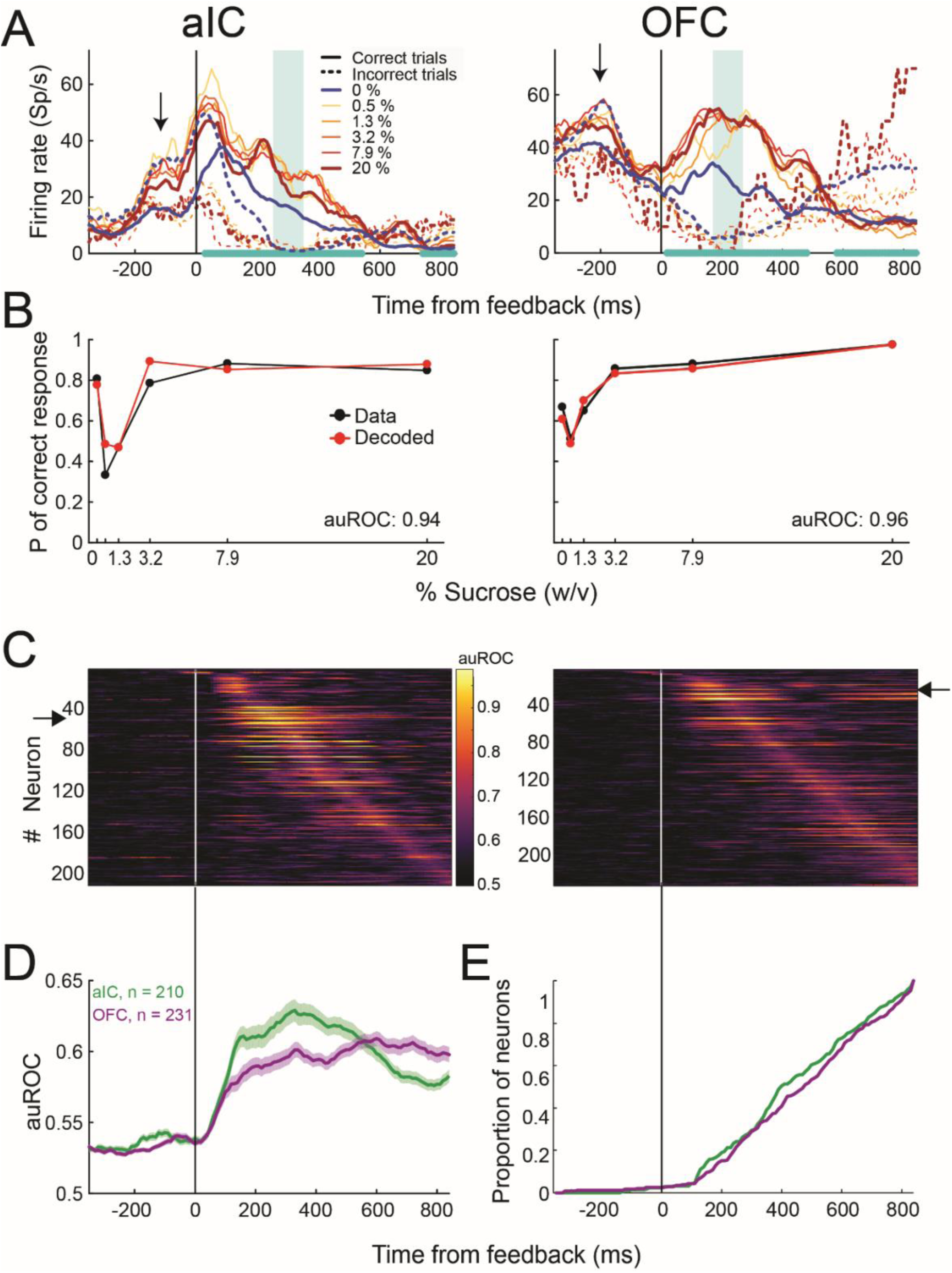
Neurons in aIC and OFC rapidly encode the outcome of the categorical decision. (**A**) The mean firing rate of one aIC (left) and one OFC (right) neuron is selective to the reward outcomes. The activity is aligned to the feedback time and segregated by stimulus condition and outcome (see inset). The cyan asterisks on the abscissa indicate the time windows with significant encoding according to a permutation test (10,000 permutations; see methods). The cyan rectangle marks the time window with the maximum significant area under the receiver operating characteristic curve [auROC] value. (**B**) Actual and decoded probabilities of correct response as a function of the stimuli for the aIC (left) and OFC (right) neurons are shown in (**A**). Actual probabilities were calculated from the number of correct trials in the experimental session where neurons in (**A**) were recorded. Decoded probabilities were calculated from the activity periods shaded in gray in the panels of (**A**) by classifying each trial as correct or incorrect, whether the neural activity surpassed the firing rate threshold that maximized the number of correctly classified trials (see Methods). **(C)** auROC values per time bin of the population of aIC (left) and OFC (right) neurons significantly encode the outcome. Neurons were ordered according to the time of maximum auROC value. Red arrows indicate the position of the neurons depicted in(**A**). (**D**) Mean (+ SEM) across time of the auROC values of the neural populations shown in (**C**). **(E)** Cumulative proportion of neurons from the populations shown in **(C)** that reached their peak auROC values across time bins. Inset shows the corresponding cumulative distributions of the times of the first lick in the lateral port. Same conventions as in **Figure 2**.

Interestingly, both neurons exhibited a diminished activity peak when the rats received water as a reward in the water choice spout compared to when they received it in the sucrose choice spout (blue vs. red solid lines, **Figure 5A**). Even though the reward had the same quality and volume in both spouts (three drops of water), the neural responses were slightly different. These results suggest that some cortical neurons could also discriminate between water as a reward depending on the spatial location (lateral spouts) from which they obtained the reward. The cause remains unclear, but one possibility is a subtle difference in drinking position or that this activity, besides encoding reward delivery, also encodes spatial location. Notably, this phenomenon has been previously observed.^30^

Given that the rats licked more the response ports in correct versus incorrect trials (**Figure S6A**), one possibility is that the aIC and OFC neural activity encoded the licking behavior of the rats instead of the trial’s outcome.^7^ We discarded this possibility by training an SVM to decode the outcome from the spiking activity or the rat’s licking behavior. We found that the licking-related decoding accuracy drops to chance (0.5) periodically, but the decoding accuracy of the neural activity remains high during the post-feedback period (**Figure S6B**). This observation suggests that the aIC and OFC neural activity could encode the trial’s outcome in addition to the rat’s licking rate.^7^

Figure 5B shows the probability of correct responses as a function of stimulus concentration calculated from the subjects’ behavioral performance (black) and decoded from the neural activity indicated by the cyan rectangle in panel **A**. Both the neurometric curves and the high auROC values suggest that the activity of these neurons is highly related to the outcome. In the aIC, 81 (38.57%) neurons exhibited preferred activity for incorrect trials (reward omission), 86 (40.95%) neurons showed higher activity for correct trials (reward delivery), and 43 (20.48%) neurons showed a shift in their response patterns (**Figure S5B**, lower panel left). In the OFC, 105 (45.45%) neurons showed higher activity in incorrect trials, 88 (38.1%) neurons had higher activity during correct trials, and 38 (16.45%) shifted their response patterns (**Figure S5B**, lower panel right). In each taste cortex, we recorded a similar proportion of neurons selective to reward omission or reward delivery, suggesting that rats could use both outcomes equally to solve the task.

Next, we explored sequential outcome encoding. The aIC and OFC outcome encodings were sorted by their peak auROCs and plotted in Figure 5C. These graphs show that the encoding of the outcome is less sequentially structured than stimulus and choice encoding (see Figures 2C **and 4C**, respectively) in the aIC and OFC. The SqI, PE, and TS indexes were 0.475, 0.983, and 0.23 for the aIC and 0.467, 0.986, and 0.221 for the OFC. TS suffered the largest diminution, suggesting that the outcome was encoded more extendedly than stimulus and choice. Conversely, we found that outcome encoding sequences were highly reproducible between splits of different trials (aIC, MSE between splits: 0.003, p < 0.001; OFC, MSE between splits: 0.002, p < 0.001). Figures 5D and **5E** show the averaged auROC of outcome-encoding cells during the outcome epoch and the cumulative distributions of the proportion of neurons that reached their peak auROC values across the outcome epoch, respectively. We observed a rapid encoding (after 50 ms) of outcome (reward delivery vs. omission) in both gustatory cortices. We found no significant differences in the temporal recruitment of aIC and OFC outcome-encoding cells (Figure 5E; Kolmogorov-Smirnov, p = 0.23), suggesting parallel recruitment and a brain-wide representation of outcome^27^ across gustatory cortices.^7^

### All relevant task variables are reflected in the principal components of aIC and OFC population activity

The previous analyses show that neural subpopulations in the aIC and OFC carry information related to the main events in the task. The weight of this information in the overall neural activity of the aIC and OFC remained unclear. Therefore, we performed a Principal Component Analysis (PCA) on the activity of all recorded neurons (including those not previously included in the analysis) in each cortical area and searched for the Principal Components (PCs) that explained the most variability in the data and how they were related to the task variables. We computed the PCs employing the neural activity averaged over the different trials of the relevant task conditions (see below and Methods). Consequently, the neural trajectories reflect mean trends and exclude variability across trials.

The upper panels in Figure 6A show the main PCs calculated from the activity of 360 aIC (left panel) and 441 OFC neurons (right panel) from 50 ms before to 800 ms after stimulus delivery. These PCs explain 30.2% and 26.9% of the total variance in the neural activity of each taste cortex. In the figure, the different colors indicate the six stimulus conditions. In both cortical areas, the trajectories diverged after stimulus delivery, reflecting the stimulus’s identity. For aIC, the divergence began and accumulated soon after stimulus delivery. In contrast, the divergence started later and was less gradual for the OFC. Remarkably, these observations coincide with the time series of average auROCs and the cumulative latency distributions shown in Figures 2D and **2E**. In addition, in the aIC, the population trajectories of the sucrose concentrations appear to cluster together, which does not occur in the water condition.

**Figure 6.**
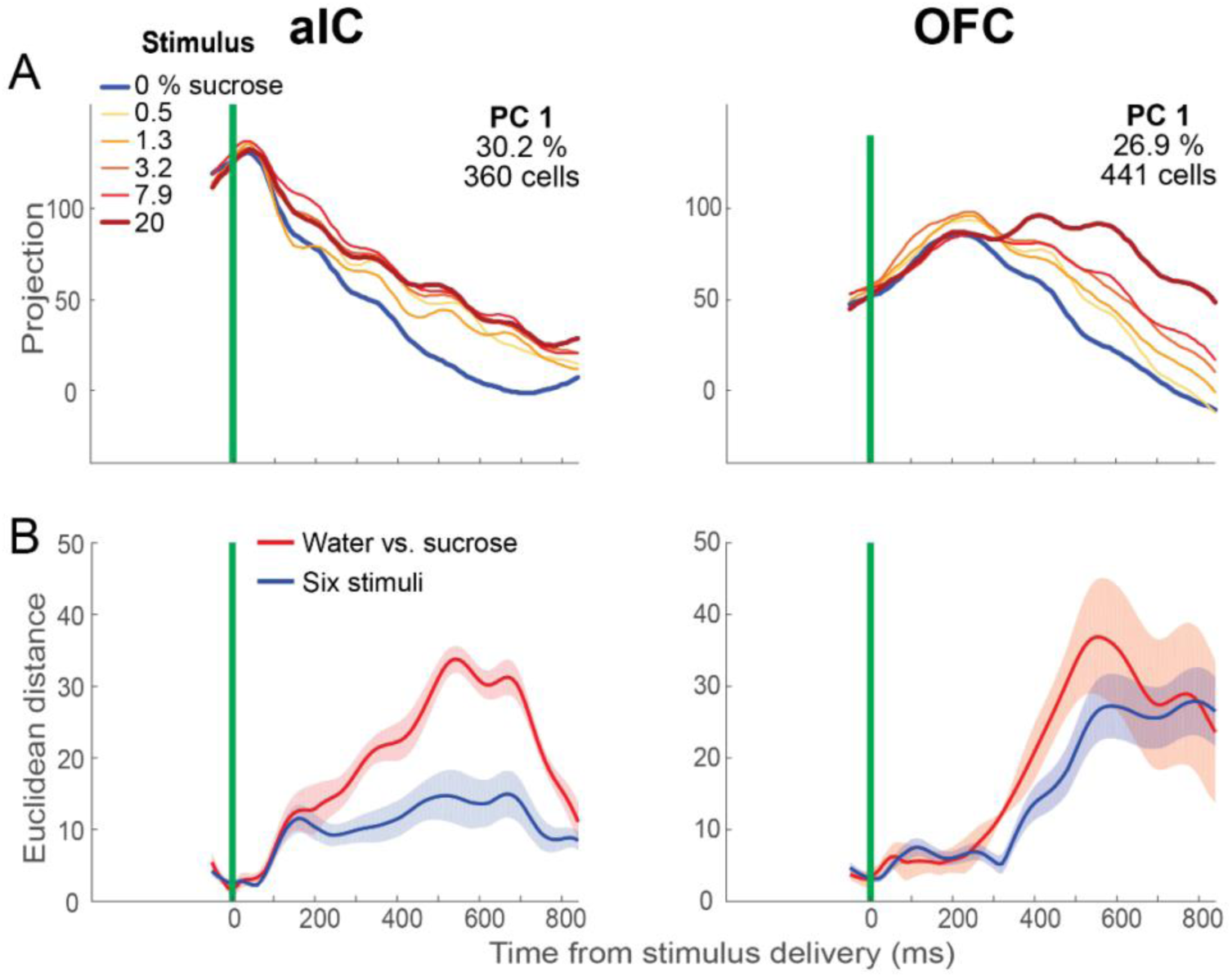
Principal components reflect the difference in stimulus selectivity observed in aIC and OFC. (**A**) Principal components (PCs) calculated from the activity of aIC (left panel) and OFC neurons (right panel). The projections are aligned to the time of stimulus delivery (green line) and are segregated by stimulus condition (the six sucrose concentrations). **(B)** Mean (± SE) Euclidean distances computed in each time bin from the projections in (**A**). The water vs. sucrose distances were calculated from all the possible water-sucrose condition pairs (five pairs). The six stimulus distances were calculated from all the stimulus pairs (15 pairs). Compare with the temporal dynamics of decoding accuracy in **Figures 3A and 3C**.

On the contrary, in the OFC, all the trajectories are more concentration-dependent in the population-state space, with water and 20% sucrose in opposite sites. To quantify this, we calculated binned Euclidean distances between the trajectories of pairs of stimulus conditions for each area. We distinguished two types of distances: the distance between the water and any sucrose conditions (water-sucrose) and the distances between all stimulus conditions (six stimuli). Figure 6B shows the mean ± SEM of these distances. In the aIC, both distances showed an early increase between 100 and 150 ms after stimulus delivery, but soon after, the water-sucrose distance became dominant over information-discriminating sucrose concentrations. In contrast, in the OFC, the PCA trajectory exhibited a delayed onset (250 ms) for discriminating stimulus information. Still, both distances (water vs. sucrose or all six stimuli) were similar over the period shown, corroborating that the OFC trajectories were more dispersed and concentration-dependent than in the aIC at the population neural-state space. Our results revealed that the aIC discriminated between taste stimuli earlier than the OFC. However, the OFC subsequently encoded more information about sucrose concentrations (perhaps reflecting its hedonic value) than the aIC.

The panels of Figure 7 depict the projections of three PCs calculated from the activity of 236 aIC (upper panels) and 191 OFC neurons (lower panels), aligned to the time of the trial’s feedback (the moment of reward delivery or omission). In this case, the activity was segregated into twelve conditions, resulting from combining the six stimuli and the two outcomes (correct or incorrect trials). Note that both the choice and the outcome are reflected in the divergence of trajectories. For example, for aIC (upper panels in Figure 7), PC 1, which explains 18.5% of the total variance, shows trajectories that segregate completely into correct and incorrect trials soon after feedback (water reward delivery or omission). In contrast, for the same cortical area, the trajectories of PC 2 are first segregated according to the rat’s decision (water or sucrose responses, see arrow), but they reconfigured soon after feedback, so the component now reflects the outcome (i.e., whether the trial was rewarded). Note that the reconfiguration is associated mainly with a ‘switch’ in the trajectory of the water condition. This PC is similar to the activity pattern of the single neuron depicted in Figure 5A. PC 5 from the aIC is also related to the choice of the rats (see arrow). For the OFC (see lower panels in Figure 7), PCs 1 (13.6%), 3 (7.8%), and 4 (6.1%) are shown. PC 1 and PC 3 trajectories were segregated according to correct and incorrect trials late or soon after feedback. In contrast, trajectories in PC 4 segregate according to the rat’s choices (see arrow) in the time preceding the feedback. Note that the categorical choices are reflected in the OFC earlier than in the aIC (compare PC 4 from the OFC to PCs 2 and 5 from the aIC).

**Figure 7.**
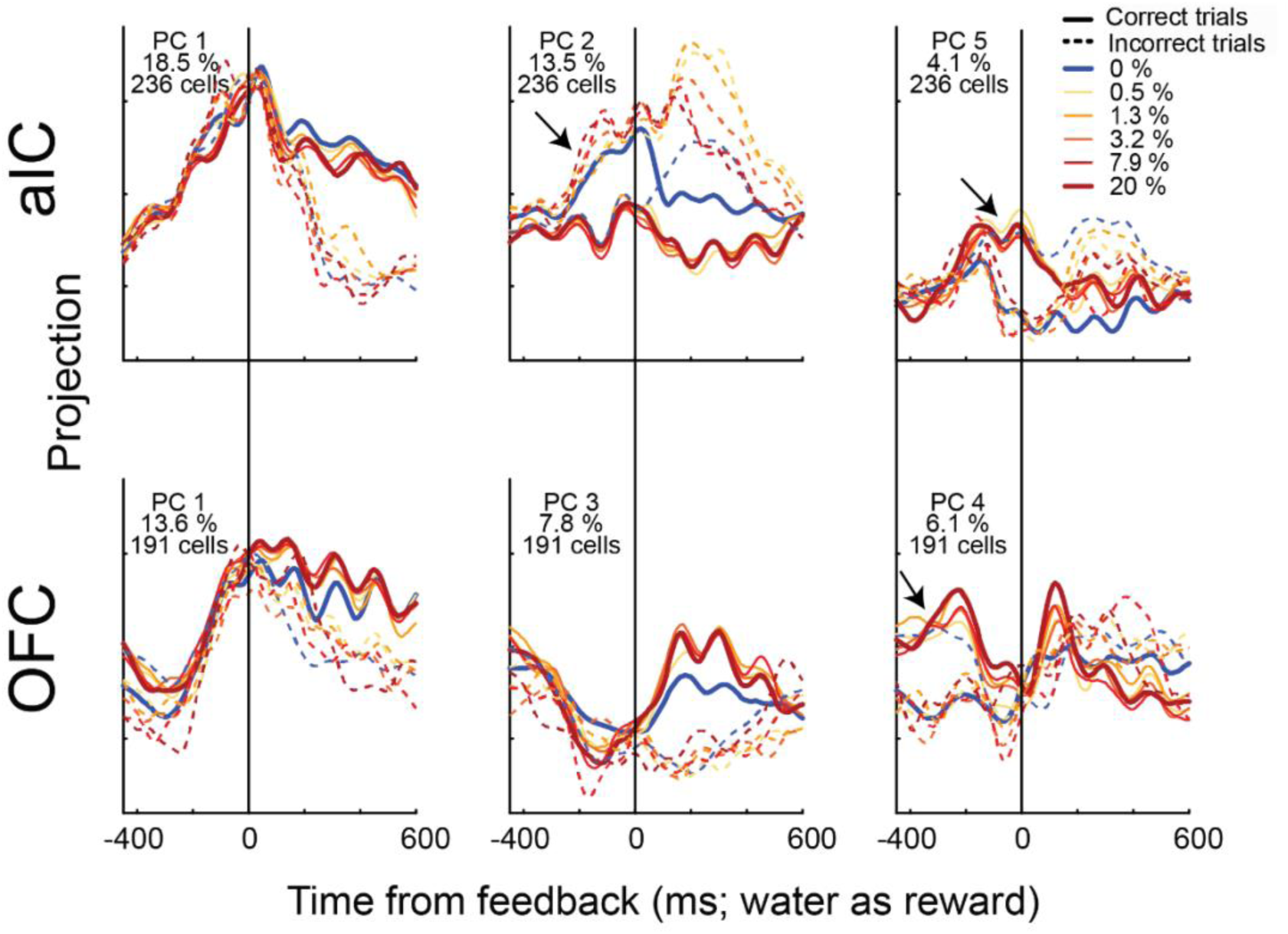
The main task variables are reflected in the principal components of the aIC and OFC neural population activity. PCs computed with the activity of 236 aIC (upper panels) and 191 OFC (lower panels) neurons aligned to the feedback time (t = 0, black vertical lines) and segregated by stimulus condition and outcome (correct or incorrect trials) as indicated in the inset in the upper right panel. Same conventions as in **Figure 6**. The arrows indicate moments when the segregation of the trajectories reflected the rat’s choices. Note that the OFC carries information about choice earlier than the aIC. Divergence of PCA trajectories after time 0 s reflexes the reward outcomes.

From the PCA, we can conclude that stimulus, choice, and outcome are the primary information encoded in the aIC and OFC while performing the sucrose categorization task. Second, whereas stimulus-related activity emerges earlier and more gradually in the aIC than in the OFC, choice-related activity emerges earlier in the OFC than in the aIC, as suggested by the single-cell analyses. Third, an essential component in aIC switches between choice and outcome representation by dynamically reconfiguring after the trial’s feedback. This is consistent with the similar pattern observed in the activity of some single aIC neurons. Collectively, PCA population analysis recapitulated most, if not all, the encoding properties observed in single neurons.

## Discussion

Our study investigated how the brain categorizes taste. We trained rats to discriminate between water and sucrose solutions in a novel task involving sampling a 10 µl drop and indicating the presence or absence of sucrose. This approach, utilizing stimuli near the detection threshold, allowed us to probe the subjective perception of taste in the subjects. As expected, rats easily categorized suprathreshold sucrose concentrations (3.2%, 7.9%, and 20%). However, near-threshold concentrations (0.5% and 1.3%) proved challenging, often being misidentified as water. Electrophysiological recordings revealed that neurons in the aIC and OFC primarily focused on differentiating water from sucrose, rather than encoding specific sweetness levels. Interestingly, the aIC accomplished this categorization faster (within 200 ms) than the OFC (over 200 ms). Contrary to observations in other species, where stimulus and choice encoding transition across the brain hierarchy, both aIC and OFC neurons in rats encoded both stimulus and choice. For example, in monkeys detecting somatosensory stimuli, for example, there is a transition from stimulus to choice encoding across the brain hierarchy, where a parameterized representation of the stimulus is found in the neural activity of the primary sensory cortices, and the neural activity in frontal areas is more related to encoding the subject’s decision.^31–34^ In rats, we found that both regions maintained similar encoding of choices throughout the experiment. Additionally, choice information emerged earlier in the OFC, which is consistent with our previous research.^7^ These results suggest a dynamic interplay between the aIC and OFC, with sensory information initially processed in the aIC and subsequently integrated with decision variables from the OFC. This distributed sensorimotor transformation framework elucidates how taste acquires behavioral meaning. A particularly striking finding was the sequential encoding of taste and choice information in gustatory cortices, resembling a “relay race” of encoding cells. This dynamic encoding challenges the traditional view of static taste representations, suggesting that taste information is continuously updated throughout the decision-making process. This study reveals the existence of a “dynamic code”^35^ and neural sequences^36^ within the gustatory system, corroborated by single-cell recordings, population decoding, and PCA. Furthermore, both the aIC and OFC rapidly encoded reward outcomes (correct/incorrect choices), further underscoring their pivotal roles in taste perception, decision-making, and quickly detecting reward outcomes.

Our previous research with rats performing sucrose discrimination and generalization tasks revealed that aIC and OFC neurons encoded various aspects of behavior, encompassing stimulus intensity (perceived sucrose concentration), choices made, movement direction, and reward outcomes.^7^ Remarkably, our current study, employing a novel sucrose categorization task, yielded similar findings. Across both tasks, we observed a consistent proportion of aIC and OFC cells encoding stimulus, choice, and outcome information. This consistency suggests a potential overarching encoding principle, transcending specific task details, and focusing on the general trial structure and decision-making processes (policies) inherent in these behavioral paradigms. However, several key questions remain. First, what is the precision of the neural representation of sucrose intensity? Can neural activity accurately differentiate between different sucrose concentrations? Second, what is the behavioral significance of these theoretical parametric representations of intensity? Lastly, the temporal encoding patterns within individual neurons of the gustatory cortices and their representation in low-dimensional population activity during decision-making remain unexplored. To address these questions, we employed time-resolved decoding techniques and PCA to analyze the activity of aIC and OFC neurons in rats performing a new sucrose categorization task.

Previous research identified cortical neurons exhibiting concentration-dependent responses to sucrose, glucose, or mixtures (in monkeys^10,12,37^ and rats^9,11^ passively exposed to various tastants). Anticipating robust stimulus intensity encoding in our experiment, our findings instead revealed that aIC and OFC stimulus-encoding cells primarily distinguished water from sucrose (quality-based encoding), rather than encoding sucrose concentration variations. While some individual neurons and population-level PC trajectories showed modulation across the six stimuli (Figures 2A**, 2B, and 6A**), our population trial-by-trial SVM decoding of the sucrose concentrations was relatively poor compared to water vs. sucrose discrimination (Figure 3 **and Figure S3**). This weak encoding likely stems from overlapping firing rates across stimulus conditions and trial-to-trial neural response variability (Figure 2B).

Interestingly, these neural encoding properties mirrored the rats’ behavior. Psychophysical performance suggests rats segregated stimuli into three groups: high sucrose concentrations (reliably categorized as ’sucrose’), 0% sucrose (reliably categorized as ’water’), and an intermediate group (near-chance categorization) (Figure 1B). This behavioral pattern, suggesting robust encoding of extreme stimuli with a blurry distinction in the middle, aligns well with neural decoding and PC trajectories, particularly in the OFC (Figure 3B**, right panel;** Figure 6A**, right panel**).

Several factors may explain the discrepancies between our findings and previous studies proposing a parametric representation of tastant concentrations. First, most prior studies relied solely on correlation or regression analyses of single-neuron activity, without attempting to decode the different stimuli. Our study, in contrast, employed decoding techniques to directly assess the neural representation of varying tastant concentrations. Previous studies identified a relatively small proportion of neurons exhibiting significant correlations/regressions between tastant concentration and neural firing rate (ranging from 0.6% to 6.45% of recorded neurons). Some studies even reported findings based on a very limited number of neurons (n=3), without quantifying the overall responsiveness.^12^ In contrast to our work, previous studies did not directly compare the relative strength of different encoding schemas (e.g., quality-based vs. intensity-based). This may have obscured the dominance of quality-based encoding observed in our study.

Our findings further confirm the sparse and distributed nature of taste coding, with only a small fraction of neurons carrying information about taste stimuli.^7,28^ Consistent with our previous work,^7^ we identified a limited population of cortical sensory neurons (<18%) exhibiting linear responses to sucrose concentration changes, both during increases (reflecting intensity changes) and decreases (potentially linked to osmotic pressure^38,39^ or bicarbonate ion in saliva^40^). This may explain the robust responses to distilled water in our categorization task, potentially overshadowing the sparse monotonic responses to sucrose concentration. These water-evoked responses further support the concept of water as an independent taste quality.^41^

Moreover, our data reveal that water-evoked responses, particularly when water serves as a reward (in lateral spouts), are multifaceted and cannot be solely explained by oral, osmotic, or purely gustatory factors. This is evidenced by subtle differences in neural activity between taste cortices depending on whether rats drank water from the left or right spout, suggesting that reward expectation,^42^ body position, or even ipsilateral or contralateral tongue movements (either left or right spouts^43^) may modulate taste-evoked responses.

Our data also showed that water-evoked responses, mainly when water was used as a reward (in lateral spouts), are more complex and could not be merely explained by oral, osmotic, or pure taste factors. This is because neural activity in both taste cortices was slightly different if rats drank water from the left or right spout,^30^ suggesting that reward expectation,^42^ differences in body position, or probably ipsilateral or contralateral tongue movements (either left or right spouts^43^) modulate taste-evoked responses.

In an active decision-making context, our study observed a general ramp-up activation pattern in the rat’s brain, reflecting a sensorimotor transformation from stimulus perception to decision-making.^27,44^ Sampling a stimulus (a quantal drop of taste molecules) triggers a neuronal avalanche, with increasing numbers of neurons participating in encoding choice, anticipation, and reward outcome. This neuronal recruitment peaks and then subsides, suggesting a brain-wide phenomenon observed across multiple cortical and subcortical regions, underscoring its importance for active sensory processing and decision-making.^7,13,27,44–46^ Crucially, our results demonstrate that the aIC and OFC are not merely passive primary or secondary sensory cortices, aligning with previous observations.^7,13,15,47^ These taste cortices actively participate and interact throughout the entire behavioral response, from initial taste perception to taste-guided decisions, employing a dynamic coding strategy in the form of sequential encoding activity.

A key discovery was the sequential and dynamic representation of taste and choice information in both the aIC and OFC. Distinct neurons encoded information at different times, forming reproducible “encoding sequences” across trial subsets. This phenomenon may appear similar to the cell assembly sequences described in many previous studies. In cell assembly sequences, neurons activate sequentially, creating reproducible patterns across trials under the same experimental conditions. These sequences are characterized by a unique combination of active neurons at each time point, allowing for decoding temporal information or information that systematically develops over time.^16,48,49^ Our study, however, revealed sequences of information encoding, not neural activation. The activity of each neuron was selective to specific task variables, and different neurons maximally encoded such information at various time points, generating a sequence of information maintenance. It is important to note that in these encoding sequences, the activation periods are not necessarily the same as the encoding periods (see, for example, the first peak of neural activity depicted in Figure 4A). Similar sequences have been observed, for example, in the encoding of choice in the parietal cortex of rats during navigation decision tasks.^36^ In their case, sequences were proposed to maintain the information for working memory and movement planning during navigation. Our study demonstrates that these sequences are not limited to navigation contexts. Another hallmark of dynamic encoding is that individual neurons change their tuning patterns.^19^ We found that some neurons even inverted their mixed-selectivity pattern. For instance, some choice-selective neurons initially showed higher activity for the water response trials but later in the trial exhibited a higher response for the sucrose response trials (**Figure S5A**, middle panels). Some outcome-selective neurons also demonstrated this encoding switching pattern (**Figure S5B**, middle panels).

A more complex switching pattern was observed in some aIC neurons that transitioned from choice to outcome encoding (Figure 5A). Interestingly, this change was systematic. Before feedback, activity during the correct water responses was more similar to that during the incorrect water responses, and activity during the correct sucrose responses was more similar to that during the incorrect sucrose responses, thus reflecting the animal’s response. After feedback, only the incorrect and correct water responses changed, so the overall activity now reflected the trial’s outcome (Figure 5A). This effect was robust enough to be seen at the population level in the PCA (see PC 2 in Figure 7, upper panel).

The relevance and neural underpinnings of dynamic coding are still under debate,^35^ but dynamic coding often emerges in complex tasks with multiple stimuli.^19^ A recent modeling study found that the emergence of dynamic codes depends on the temporal complexity of the task, the variability of delay durations, the strength of neuron coupling, prior training in another task, and the presence or absence of task-irrelevant, motion-related dynamic cues.^50^ Studies comparing tasks with diverse complexities and structures will shed more light on these questions.

### Limitations of the study

Here, a caution note is required; as we analyzed the activity of neural pseudo-populations, the encoding of the different stimuli could have been lost because of the variable duration of the trials.^51^ A related alternative is that stimulus encoding might not reside in the firing rate alone. Previous studies proposed that spike timing, besides firing rate, may play a local role in taste processing.^7,13,52^ Our data did not rule out that spike timing could contain additional information to distinguish specific concentrations. Simultaneous recordings, unsupervised time-warping methods, and spike-time-based analysis are required to test these possibilities. Alternatively, aIC and OFC activity perhaps prioritized categorizing stimulus information over specific taste concentration because of the behavioral context.^53^ To solve our categorization task, rats only needed to distinguish water from sucrose, regardless of the concentration; thus, the intensity attribute is not entirely relevant to solving this task. Perhaps in a sweetness intensity discrimination task,^7^ gustatory cortices would allocate more attention to this sensory attribute. If confirmed, this would further suggest that gustatory cortices activity is context-dependent and adapts to the demands of the behavioral task.^54^

## Conclusion

In summary, in a new paradigm of categorizing different sucrose concentrations, the firing rate of aIC and OFC neurons did not maintain a sharp, parametric representation of the concentrations but rather a robust, gross distinction between sucrose and non-sucrose solutions. Furthermore, task-relevant information was not maintained by long-lasting firing rate codes but by sequences of information encoding, in which the firing rate of different neurons carries the relevant information at distinct times and for brief periods during the trial (see Graphical Abstract). aIC and OFC neural populations also exhibited changing patterns of mixed-information encoding, another hallmark of dynamic encoding. The presence of sequential and dynamic encoding in the gustatory system provides valuable insights into how gustatory cortices transform taste information into neural signals that ultimately guide taste-based decisions.

## Acknowledgments

We thank Mario Gil Moreno for building multielectrode arrays and Victor Manuel García Gomez, and Fabiola Hernández Olvera for their invaluable animal care. We especially want to thank Francisco Zepeda Arias and Silvia Mejía Ortiz for their invaluable help in training the subjects.

## Author contributions

E.F. and R.G. conceived the project. E.F. conducted experiments and performed data curation. G.M. performed code implementation and formal analysis. R.G. supervised the project and provided resources. R.G. and H.M. supervised the analysis. All authors discussed the results. G.M. wrote the original draft. All authors reviewed, edited, and approved the final manuscript.

## STAR★Methods

### Resource availability

#### Lead contact

Further information and resource requests should be directed to, and will be fulfilled by, the lead contact, Dr. Ranier Gutierrez.

#### Materials availability

This study did not generate new unique reagents.

#### Data and code availability

- All original code has been deposited and is publicly available as of the date of publication. DOIs are listed in the key resources table.

#### Experimental model and subject details

Male Sprague-Dawley rats (n = 11) weighing 300–320g at the beginning of the experiment were trained. The subjects were individually housed in cages in a temperature-controlled (22 ± 1°C) room, with a 12:12 h light-dark cycle (lights on at 0700 h) and ad libitum access to food (PicoLab Rodent Diet 20, St. Louis, MO, USA). Experiments were conducted at the same time within the light-dark cycle (1400 to 1900 h). After each experiment, the rats were given free access to water for 30 min. The CINVESTAV Institutional Animal Care and Use Committee approved all the experimental procedures.

### Method details

#### Apparatus

Standard operant conditioning chambers were employed (30.5 x 24.1 x 21.0 cm, Med Associates Inc., VT, USA). The front panel of each chamber was equipped with one central and two lateral V-shaped lick spouts with a photobeam sensor to register individual licks (Med Associates Inc., VT, USA). The central spout comprised a bundle of six blunted needles (20 gauge) glued at the tip of a stainless-steel sipper tube, while each lateral spout comprised one needle. Each needle was independently connected to pressurized liquid reservoirs using a silicone tube, resulting in an eight-channel fluid delivery system.^7^ By turning on and off independent solenoid valves (Parker, Ohio, USA), the system allowed the delivery of controlled volumes of liquid in each spout. One of the six needles in the central spout delivered the stimulus concentrations, while the lateral spouts delivered the water reward. On the rear panel was an ambiance white noise amplifier with a speaker that was turned on during all sessions. Chambers were enclosed in a ventilated sound-attenuating cubicle. Experimental events were controlled and registered by a computer via a Med Associates interface (Med Associates Inc., VT, USA).

#### Stimuli

Six solutions (0, 0.5, 1.3, 3.2, 7.9, and 20 w/v%) of sucrose (reagent-grade, Sigma-Aldrich, Mexico) were diluted in distilled water. All sucrose concentrations were prepared every other day, maintained under refrigeration, and used at room temperature during the experiments.

#### Sucrose categorization task

Rats were trained in a two-alternative forced-choice task to report the presence or absence of sucrose in a 10 µL stimulus drop. The trial structure is as follows: rats were required to reach and dry-lick the stimulus spout two or three times to receive a 10 µL liquid drop (stimulus delivery) (Figure 1A). The stimulus could be one of six sucrose concentrations: 0, 0.5, 1.3, 3.2, 7.9, and 20%. Subsequently, the subject had to move and dry-lick the lateral response Spout associated with the stimulus received: Water stimulus → Left response spout; Sucrose stimulus → Right response spout. These conditions were counterbalanced; thus, we will refer to response spouts as water and sucrose response spouts. If the rat licked the correct response spout, the subsequent three licks resulted in the delivery of one drop of water per lick as a reward; otherwise, for error trials, the lights went off and on, and no reward was provided. The moment of reward delivery or omission is designated as the time of feedback. Water (0% sucrose) was used as the stimulus in half of the trials per experimental session, while the remaining sucrose concentrations were homogeneously distributed in the other half. The learning criterion was 80% correct responses during four consecutive sessions.

#### Electrode implantation

Once animals achieved the learning criterion, custom-made, 4 x 4 tungsten wire electrode arrays were implanted in the left hemisphere from aIC (n = 6) or OFC (n = 5) under ketamine/xylazine anesthesia (70 mg/kg/20 mg/kg, i.p.).^14^ Implantation coordinates were as follows: aIC, +1.6 to +2.3 mm AP, +5.2 mm ML from bregma, and -4.6 to -4.7 mm DV from the dura; OFC, +3.5 mm AP, +3.2 mm ML from bregma, and -4.4 mm DV from the dura. After surgery, rats were given intraperitoneal enrofloxacin (0.4 ml/kg) and ketoprofen (45 mg/kg) for three days and allowed one week of rest for recovery.

#### Electrophysiology

Extracellular voltage signals acquired by a MAP system (Plexon, Dallas, TX) were amplified x1 (analog headstage, Plexon HST/16o25-GEN2-18P-2GP-G1) and x1000 and sampled at 40 kHz. Raw signals were band-pass filtered (154 Hz-8.8 kHz) and digitalized (12-bit resolution). The action potentials were isolated using voltage-time threshold windows and PCA in real time. Waveforms were assigned to the same unit if the refractory period (1 ms) was not violated and whether a neat delimited ellipsoid cloud was on the first three PCs. Additional offline sorting was performed (Offline Sorter, Plexon, Dallas, TX), and only single units with a signal-to-noise ratio of 3:1 were analyzed.^7,14^

#### Histology

After the experiments were completed, subjects were administered an overdose of pentobarbital sodium (150 kg/mg, i.p.) and perfused transcardially with PBS (1x) followed by 4% PFA. Brains were removed, stored for 24 h in 4% PFA, and changed to a 30 v/v% sucrose/PBS solution. Coronal, 40-micron sections stained with Cresyl violet corroborated the recording locations.^7^

### Quantification and statistical analysis

Subroutines written in Matlab (Matworks version 2016b) and the SPSS statistical package (version 29.0.2.0, SPSS Inc., Chicago, IL) were used for the statistical analyses. The level of statistical significance to reject the null hypothesis was α = 0.05 unless stated otherwise.

#### Behavior

The psychometric curves for each session were built as the probability of selecting the sucrose-response spout (‘p of sucrose response’) for each stimulus concentration. Power functions (f_(x)_ = ax^b^) were fitted to these data, and the stimulus value at 0.5 probability of responding ‘sucrose’ was computed as the absolute threshold. The mean values of the psychometric curves oscillate between 0 and 1, but no stimulus concentration leads to perfect categorization, not even the “obvious/easiest” concentrations of 0 and 20% sucrose. To corroborate that stimulus presentations were balanced within sessions, we quantified and compared the proportion of trials in which the subject received water or sucrose stimulus by paired samples and a one-tailed t-test. We also characterized the subject’s responses and their associated outcome distribution: we quantified and compared, by a paired sample, one-tailed t-test, the proportion of trials in which the subject reported perceiving the stimulus as water or sucrose (water response vs. sucrose response) and the proportion of these trials that were rewarded. Finally, we evaluated the effect of stimulus concentration over the time of licking in the stimulus and response spouts and the time to move from the stimulus to the response spout. Specifically, for each stimulus concentration, we computed the time elapsed between stimulus delivery and the last lick emitted in the Stimulus spout (central spout licking), the time between the first and last Response spout lick (lateral spout licking), the time between the last stimulus spout lick and first lateral spout lick (movement time).

#### Encoding of the task’s variables

For each neuron and trial, firing rates were computed in 100-ms size windows moving in 10-ms steps. These data were employed to identify and quantify single neurons that significantly encoded Stimulus, Choice, and Outcome. The difference in the proportion of cells encoding each task variable among aIC and OFC was determined by a two-proportion Z-test. A detailed description of the analysis performed is described below.

#### Single neuron encoding of the stimulus quality

To determine the relationship between neural activity and the task’s parameters, we employed auROC analyses on the activity of each neuron. For stimulus encoding, we constructed, for each 100 ms size window, two firing rate distributions, one with the activity of each neuron during water or Sucrose stimulus trials; all sucrose concentrations are embedded in the sucrose category. auROC measures the overlap between the two distributions, with values ranging from 0 to 1, where the edge values 0 and 1 signal complete segregation of the distributions, and 0.5 indicates complete overlap.^50^ When analyzing only correct trials in these types of tasks, a trial-by-trial correlation between Stimulus and Response could emerge. Consequently, differentiating the neural activity associated with the Stimulus from the activity related to the subject’s response is a common problem. We addressed this problem by computing two auROC values from the same trials for each neuron and time bin. The firing rate distributions were arranged according to the presented stimulus, or the responses reported by the subjects. Then, we compared both auROC values and considered putative Stimulus-encoding the bins with the Stimulus-auROC higher than 0.6 (or lower than 0.4) and higher (lower) than the corresponding Response-auROC. auROC values of 0.4 and 0.6 roughly corresponded to the values at 0.25 and 0.95 in the distribution of auROC values from all the time bins and cells in this analysis. Ten thousand permutations of the Stimulus labels were employed to build null distributions per time bin to identify the bins with stimulus-auROCs above chance (p ≤ 0.05). Neurons with at least five consecutive significant bins were considered stimulus-encoding cells. For the display of single-neuron activity in the figures (Figure 2A), we employed asterisks to indicate the time windows with significant encoding and a cyan rectangle to mark the time window with the maximum significant area under the receiver operating characteristic curve [auROC] value. We also computed the mean auROC value across time for the population of encoding neurons in aIC and OFC (Figure 2D). Finally, to compare the recruiting times of neuron encoding in aIC and OFC, we computed the cumulative proportion to time of the auROC peak values of the encoding populations and compared them by a two-sample Kolmogorov-Smirnov test (Figure 2E).

#### Single-neuron encoding of the subject’s Response

To identify cells encoding the perceptual choice of the subjects, we performed an auROC analysis of firing rate distributions associated with Water or sucrose response trials. This index reflects the proportion of behavioral choices predicted from single-neuron activity on a trial-by-trial basis. By including correct and error trials in the analysis, the index eliminates the trial-by-trial correlation between the presented stimulus and the subject’s choices when analyzing only correct trials ^50,51^. For each neuron and time bin, we constructed two firing rate distributions, one containing the firing rates of the trials where the rats chose the water response and another with the firing rates of the trials where the rats chose the sucrose response. Trials of any stimulus condition could contribute to one or another distribution. From these distributions, the auROC curves were calculated. Neurons with five consecutive bins with a choice probability larger than 0.6 or lower than 0.4 and surpassing a two-tailed permutation test (10,000 permutations) were considered to significantly encode the animals’ choice. auROC values of 0.4 and 0.6 roughly corresponded to the values at 0.25 and 0.95 in the distribution of auROC values from all the time bins and cells included in this analysis. For the display of single-neuron activity in the figures (Figure 4A), we employed asterisks to indicate the time windows with significant encoding and a cyan rectangle to mark the time window with the maximum significant area under the receiver operating characteristic curve [auROC] value. We also computed the mean auROC value across time for the population of encoding neurons in aIC and OFC (Figure 4D). Finally, to compare the recruiting times of neuron encoding in aIC and OFC, we computed the cumulative proportion to time of the auROC peak values of the encoding populations and compared them by a two-sample Kolmogorov-Smirnov test (Figure 4E).

#### Single-neuron encoding of the trial’s outcome

To identify cells encoding the outcome, we calculated the outcome auROC values by comparing the firing rate distributions of rewarded vs. non-rewarded trials. Outcome-encoding neurons are determined using the same significant criterion described above. For the display of single-neuron activity in the figures (Figure 5A), we employed asterisks to indicate the time windows with significant encoding and a cyan rectangle to mark the time window with the maximum significant area under the Receiver Operating Characteristic curve (auROC) value. We also computed the mean auROC value across time for the population of encoding neurons in aIC and OFC (Figure 5D). Finally, to compare the recruiting times of neuron encoding in aIC and OFC, we computed the cumulative proportion to time of the auROC peak values of the encoding populations and compared them by a two-sample Kolmogorov-Smirnov test (Figure 5E).

#### Sequences of encoding by single neurons

Across the neural population, different neurons encoded the task’s variables at various time points. To visualize this phenomenon, we sorted neurons according to the time of the peak auROC value, which resulted in sequences of neural encoding for each cortical area and task’s variable (see Figures 2C**, 4C,** and **5C**). To quantify the degree of this sequential encoding in aIC and OFC populations, we applied the Sequentiality Index (SqI)^25^. The original index is calculated on the firing rates of the neural populations and is bounded between 0 and 1. The SqI approaches 1 when the peak times of each neuron homogeneously tile the entire duration of interest, and one neuron is active at every moment in time, with the temporal fields being non-overlapping. The index approaches 0 otherwise. The SqI is computed from the Peak Entropy (PE), which measures the entropy of the distribution of peak times across the entire neural population, and Temporal Sparsity (TS), which measures the entropy of the distribution of normalized activity in any bin. We computed PE, TS, and the Sql with the data from the matrix of sorted auROC values (only auROCs above 0.6 or below 0.4 were included in the calculus) for each task’s variable and cortical area, from zero to 800 ms after the alignment time. Used in this way, PE reflects how homogeneously the encoding times of the different neurons tile the analyzed period, with values close to one for more homogenous covering and zero otherwise; TS measures how much of the activity at each point in time can be attributed to a single neuron, with zero and one corresponding to overlapped and not-overlapped encoding periods, respectively. The SqI measures the level of sequential information encoding by the neural populations in aIC and OFC.

We performed holdout cross-validation to evaluate the reproducibility of the neural encoding sequences across trials. First, we randomly partitioned the data from each neuron into two trial subsets and ran the ROC analysis separately for each data split and neuron. We constructed two matrices of population auROC values across time (cell-by-bin size matrices). We then ordered the cells in matrix two by the indexes resulting from sorting the cells in matrix one by their latencies to peak auROC. Next, we calculated the mean-squared error (MSE; Matlab *immse* function) between the two sorted matrices to measure pattern similarity. MSEs close to zero imply that the population encoding dynamic is similar across the two data splits. A permutation test (10,000 permutations of the row indexes of matrix 2) allowed us to test the possibility that such similarity resulted from chance.

#### Support Vector Machine populational decoding

We used 8-fold cross-validated (*crossval* function, Matlab), one vs. one Support Vector Machine (SVM, *fitcecoc* function, Matlab) for decoding the Stimulus delivered in each trial from neural population activity. We trained the SVM to decode Water versus Sucrose, or the six Stimulus concentrations in 100-ms time windows, moving in 10-ms steps from -400 to 800 ms from the Stimulus delivery. We ran the SVM with linear, Gaussian, and third-order polynomial kernels. We took the classification loss from observations not used for training (1 – loss, *kfoldLoss* function, Matlab) as a measure of decoding accuracy to compare this accuracy across cortical areas and decoding schemas (Water vs. sucrose or the six concentrations). For this purpose, the SVM was trained in the two decoded schemas with the same number of neurons, trials per neuron, and trials per decoded condition. For water and sucrose stimulus decoding, 16 correct trials (8 trials per condition) were employed per SVM run. To decode the stimuli, 48 correct trials per stimulus condition were used. Homogenizing the trial number required the selection of neurons with 2 x 8 correct trials when arranged according to the binary decoding schema and 6 x 8 correct trials when arranged according to the other schema. Using the same neurons for both decoding schemas allowed direct comparisons within cortical areas. Also, note that the employed trial’s counts would have favored decoding the six concentrations. The number of neurons and trials was homogenized by random sampling with replacement across brain areas and neurons. Fifteen samplings for neurons and ten for trials were utilized for 150 samplings. Decoding accuracy is reported as the mean (± SEM) of the 150 samplings. Stimulus labels were permuted to calculate the chance of SVM performance. Permutation results are the mean (± SEM) Stimulus decoding accuracy of 150 neuron/trial samplings. The whole analysis was first performed on the activity of aIC and OFC Stimulus-encoding neurons from the Signal Detection Theory-based analysis. Next, the analysis was run on the activity of these cells and all the neurons with significant differences in a one-way ANOVA with Stimulus as the six-level factor. Neurons with significant choice encoding, according to auROC, were excluded from the analysis. To further investigate whether the performance of the decoder trained with aIC or OFC neural activity is more similar to gross discrimination between water and sucrose than to a fine discrimination of each sucrose concentration we computed confusion matrices. We counted the labels the linear-kernel SVM decoder assigned to each of the six stimuli across time bins (from 300 to 840 ms after stimulus delivery; see Figure 3A) and trial and neuron samplings. With these data, we constructed six-by-six matrices (**Figure S4**). We calculated the mean-squared error (MSE; Matlab *immse* function) between the matrices computed with aIC (**Figure S4C**) or OFC (**Figure S4D**) neural activity and theoretic fine (**Figure S4A**) or gross patterns (**Figure S4B**) to measure pattern similarity.

#### Determining the effect of licking movements

To test whether the activity of the population of outcome-encoding neurons was more related to reward delivery/omission than to the licking behavior of the rats, we compared the performance of the population neural activity or the licking behavior in the trial-by-trial decoding of the delivery or omission of reward by an SVM. If the activity of the population of outcome-encoding neurons is independent of the licking behavior of the rats, then differences between the classification accuracy of both parameters are expected^7^. First, we computed the firing rate of the outcome by encoding cells in 100 ms time windows, moving in 10-ms steps from -400 to 800 ms from the trial’s feedback. The lick rate was computed in sliding windows of the same size and steps as those used for neural activity. Only neural firing rates and licking rates from the same sessions and trials were employed. Licking times varied from trial to trial; hence, when all session trials are used for decoding, licking movements tend to ‘tile’ the post-feedback period. This could cause artificially continuous and high decoding performances when the SVM uses the licking rate. To avoid this problem, all the inter-lick intervals were normalized to 140 ms, and the spike times were warped to the normalized times. This procedure resulted in the alignment movements during licking and the warping of spike times in all the trials employed in the analysis. Next, to decode the outcome of each trial using neural population firing rates or rat licking rates, we used a linear-kernel SVM (one vs. one, 8-fold cross-validated). We took the classification loss from observations not used for training to measure the decoding accuracy for both firing rate and lick rate. The number of neurons and trials was homogenized by random sampling with replacement across brain areas and neurons (16 trials per neuron, eight from correct and eight from incorrect trials) to compare the decoding accuracy across cortical regions. Fifteen samplings for neurons and 10 for trials were employed for 150 samplings. Also, the stimulus labels were permuted to gain insight into the chance of SVM performance. Significant differences between the means of both decoding accuracy distributions were determined across time by a bootstrap test (sample with replacement; 10,000 iterations).

#### PCA and population trajectories

For each cortical area, PCAs were run over the activity of all recorded neurons averaged within the experimental conditions of interest: presented stimulus (6 conditions), animal’s choice x presented stimulus (12 conditions), or trial outcome x presented stimulus (12 conditions). For these analyses, the action potential times of each neuron in each trial were aligned to the relevant behavioral event: stimulus delivery time, time of last central spout lick, or feedback time. The activity was then convolved at 1 ms resolution, with a 20 ms width Gaussian kernel employing the *ksdensity* function in Matlab. The data were then converted to firing rate and averaged per neuron within the relevant experimental conditions. The stimulus encoding analysis included only neurons with at least five correct trials per condition. In the other analyses, we required at least two trials per condition per neuron.

The mean firing rate of all recorded neurons was arranged in a matrix of *n* x *c* with *n* = number of analyzed neurons and *c* = time bin x experimental condition. Covariance matrices were computed from these data and subsequently employed to compute PC coefficients.

Given a linear transformation of a matrix *X* into a matrix *Y*, such that each dimension of *Y* explains the variance of the original data *X* in descending order, PCA can be described as the search for matrix *P* that transforms *X* into Y as follows:

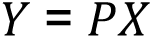

The PC coefficient matrix *P* was multiplied by the *X’* matrix to transform the neural data into the space of the original *Y*. A locally weighted smoothing function was applied to the columns of the *Y* matrix. Selected dimensions of *Y* were plotted to generate graphical one-dimensional trajectories of the population neural activity ^52^.

## References

1. Gutierrez, R., Fonseca, E., and Simon, S.A. (2020). The neuroscience of sugars in taste, gut-reward, feeding circuits, and obesity. Cell Mol Life Sci 77, 3469–3502. 10.1007/s00018-020-03458-2.

2. Barretto, R.P.J., Gillis-Smith, S., Chandrashekar, J., Yarmolinsky, D.A., Schnitzer, M.J., Ryba, N.J.P., and Zuker, C.S. (2015). The neural representation of taste quality at the periphery. Nature 517, 373–376. 10.1038/nature13873.

3. Wu, A., Dvoryanchikov, G., Pereira, E., Chaudhari, N., and Roper, S.D. (2015). Breadth of tuning in taste afferent neurons varies with stimulus strength. Nat Commun 6, 8171. 10.1038/ncomms9171.

4. Nelson, G., Hoon, M.A., Chandrashekar, J., Zhang, Y., Ryba, N.J., and Zuker, C.S. (2001). Mammalian sweet taste receptors. Cell 106, 381–390. 10.1016/s0092-8674(01)00451-2.

5. Vincis, R., and Fontanini, A. (2019). Central taste anatomy and physiology. Handb Clin Neurol 164, 187–204. 10.1016/B978-0-444-63855-7.00012-5.

6. Rolls, E.T., Yaxley, S., and Sienkiewicz, Z.J. (1990). Gustatory responses of single neurons in the caudolateral orbitofrontal cortex of the macaque monkey. Journal of Neurophysiology 64, 1055–1066. 10.1152/jn.1990.64.4.1055.

7. Fonseca, E., de Lafuente, V., Simon, S.A., and Gutierrez, R. (2018). Sucrose intensity coding and decision-making in rat gustatory cortices. eLife 7, e41152. 10.7554/eLife.41152.

8. Kogan, J.F., and Fontanini, A. (2024). Learning enhances representations of taste-guided decisions in the mouse gustatory insular cortex. Current Biology. 10.1016/j.cub.2024.03.034.

9. Maier, J.X., and Katz, D.B. (2013). Neural dynamics in response to binary taste mixtures. Journal of Neurophysiology 109, 2108–2117. 10.1152/jn.00917.2012.

10. Scott, T.R., Plata-Salaman, C.R., Smith, V.L., and Giza, B.K. (1991). Gustatory neural coding in the monkey cortex: stimulus intensity. Journal of Neurophysiology 65, 76–86. 10.1152/jn.1991.65.1.76.

11. Stapleton, J.R., Lavine, M.L., Wolpert, R.L., Nicolelis, M.A.L., and Simon, S.A. (2006). Rapid Taste Responses in the Gustatory Cortex during Licking. J Neurosci 26, 4126–4138. 10.1523/JNEUROSCI.0092-06.2006.

12. Thorpe, S.J., Rolls, E.T., and Maddison, S. (1983). The orbitofrontal cortex: neuronal activity in the behaving monkey. Exp Brain Res 49, 93–115. 10.1007/BF00235545.

13. Gutierrez, R., Simon, S.A., and Nicolelis, M.A.L. (2010). Licking-Induced Synchrony in the Taste–Reward Circuit Improves Cue Discrimination during Learning. J. Neurosci. 30, 287–303. 10.1523/JNEUROSCI.0855-09.2010.

14. Lang, L., Camera, G.L., and Fontanini, A. (2023). Temporal progression along discrete coding states during decision-making in the mouse gustatory cortex. PLOS Computational Biology 19, e1010865. 10.1371/journal.pcbi.1010865.

15. Vincis, R., Chen, K., Czarnecki, L., Chen, J., and Fontanini, A. (2020). Dynamic Representation of Taste-Related Decisions in the Gustatory Insular Cortex of Mice. Curr Biol 30, 1834–1844.e5. 10.1016/j.cub.2020.03.012.

16. Zhou, S., Masmanidis, S.C., and Buonomano, D.V. (2020). Neural Sequences as an Optimal Dynamical Regime for the Readout of Time. Neuron 108, 651–658.e5. 10.1016/j.neuron.2020.08.020.

17. Crowe, D.A., Zarco, W., Bartolo, R., and Merchant, H. (2014). Dynamic Representation of the Temporal and Sequential Structure of Rhythmic Movements in the Primate Medial Premotor Cortex. J. Neurosci. 34, 11972–11983. 10.1523/JNEUROSCI.2177-14.2014.

18. Merchant, H., Pérez, O., Bartolo, R., Méndez, J.C., Mendoza, G., Gámez, J., Yc, K., and Prado, L. (2015). Sensorimotor neural dynamics during isochronous tapping in the medial premotor cortex of the macaque. Eur J Neurosci 41, 586–602. 10.1111/ejn.12811.

19. Lundqvist, M., Herman, P., and Miller, E.K. (2018). Working Memory: Delay Activity, Yes! Persistent Activity? Maybe Not. J Neurosci 38, 7013–7019. 10.1523/JNEUROSCI.2485-17.2018.

20. Crowe, D.A., Goodwin, S.J., Blackman, R.K., Sakellaridi, S., Sponheim, S.R., MacDonald, A.W., and Chafee, M.V. (2013). Prefrontal neurons transmit signals to parietal neurons that reflect executive control of cognition. Nat Neurosci 16, 1484–1491. 10.1038/nn.3509.

21. Murray, J.D., Bernacchia, A., Roy, N.A., Constantinidis, C., Romo, R., and Wang, X.-J. (2017). Stable population coding for working memory coexists with heterogeneous neural dynamics in prefrontal cortex. Proc Natl Acad Sci U S A 114, 394–399. 10.1073/pnas.1619449114.

22. Betancourt, A., Pérez, O., Gámez, J., Mendoza, G., and Merchant, H. (2023). Amodal population clock in the primate medial premotor system for rhythmic tapping. Cell Rep 42, 113234. 10.1016/j.celrep.2023.113234.

23. Kaufman, M.T., Churchland, M.M., Ryu, S.I., and Shenoy, K.V. (2014). Cortical activity in the null space: permitting preparation without movement. Nat Neurosci 17, 440–448. 10.1038/nn.3643.

24. Perez, I.O., Villavicencio, M., Simon, S.A., and Gutierrez, R. (2013). Speed and accuracy of taste identification and palatability: impact of learning, reward expectancy, and consummatory licking. American Journal of Physiology-Regulatory, Integrative and Comparative Physiology 305, R252–R270. 10.1152/ajpregu.00492.2012.

25. Samuelsen, C.L., and Fontanini, A. (2017). Processing of Intraoral Olfactory and Gustatory Signals in the Gustatory Cortex of Awake Rats. J Neurosci 37, 244–257. 10.1523/JNEUROSCI.1926-16.2016.

26. Lara, A.H., Kennerley, S.W., and Wallis, J.D. (2009). Encoding of gustatory working memory by orbitofrontal neurons. J Neurosci 29, 765–774. 10.1523/JNEUROSCI.4637-08.2009.

27. Laboratory, I.B., Benson, B., Benson, J., Birman, D., Bonacchi, N., Carandini, M., Catarino, J.A., Chapuis, G.A., Churchland, A.K., Dan, Y., et al. (2023). A Brain-Wide Map of Neural Activity during Complex Behaviour. Preprint at bioRxiv, 10.1101/2023.07.04.547681 https://doi.org/10.1101/2023.07.04.547681.

28. Chen, K., Kogan, J., and Fontanini, A. (2020). Spatially Distributed Representation of Taste Quality in the Gustatory Insular Cortex of Behaving Mice. Current Biology. 10.1016/j.cub.2020.10.014.

29. Ben-Hur, A., Ong, C.S., Sonnenburg, S., Schölkopf, B., and Rätsch, G. (2008). Support Vector Machines and Kernels for Computational Biology. PLOS Computational Biology 4, e1000173. 10.1371/journal.pcbi.1000173.

30. MacDonald, C.J., Meck, W.H., Simon, S.A., and Nicolelis, M.A.L. (2009). Taste-guided decisions differentially engage neuronal ensembles across gustatory cortices. J Neurosci 29, 11271– 11282. 10.1523/JNEUROSCI.1033-09.2009.

31. Romo, R., Merchant, H., Zainos, A., and Hernández, A. (1997). Categorical perception of somesthetic stimuli: psychophysical measurements correlated with neuronal events in primate medial premotor cortex. Cereb Cortex 7, 317–326. 10.1093/cercor/7.4.317.

32. Romo, R., Merchant, H., Zainos, A., and Hernández, A. (1996). Categorization of somaesthetic stimuli: sensorimotor performance and neuronal activity in primary somatic sensory cortex of awake monkeys. Neuroreport 7, 1273–1279.

33. Romo, R., and de Lafuente, V. (2013). Conversion of sensory signals into perceptual decisions. Prog Neurobiol 103, 41–75. 10.1016/j.pneurobio.2012.03.007.

34. Zhou, Y., and Freedman, D.J. (2019). Posterior parietal cortex plays a causal role in perceptual and categorical decisions. Science 365, 180–185. 10.1126/science.aaw8347.

35. Stroud, J.P., Duncan, J., and Lengyel, M. (2024). The computational foundations of dynamic coding in working memory. Trends in Cognitive Sciences, S1364661324000536. 10.1016/j.tics.2024.02.011.

36. Harvey, C.D., Coen, P., and Tank, D.W. (2012). Choice-specific sequences in parietal cortex during a virtual-navigation decision task. Nature 484, 62–68. 10.1038/nature10918.

37. Rolls, E.T. (2023). The orbitofrontal cortex, food reward, body weight and obesity. Social Cognitive and Affective Neuroscience 18, nsab044. 10.1093/scan/nsab044.

38. Hanamori, T. (2001). Effects of Various Ion Transport Inhibitors on the Water Response in the Superior Laryngeal Nerve in Rats. Chemical Senses 26, 897–903. 10.1093/chemse/26.7.897.

39. Lyall, V., Heck, G.L., DeSimone, J.A., and Feldman, G.M. (1999). Effects of osmolarity on taste receptor cell size and function. American Journal of Physiology-Cell Physiology 277, C800– C813. 10.1152/ajpcell.1999.277.4.C800.

40. Zocchi, D., Wennemuth, G., and Oka, Y. (2017). The cellular mechanism for water detection in the mammalian taste system. Nat Neurosci 20, 927–933. 10.1038/nn.4575.

41. Rosen, A.M., Roussin, A.T., and Di Lorenzo, P.M. (2010). Water as an Independent Taste Modality. Front. Neurosci. 4. 10.3389/fnins.2010.00175.

42. Mazzucato, L., La Camera, G., and Fontanini, A. (2019). Expectation-induced modulation of metastable activity underlies faster coding of sensory stimuli. Nat Neurosci 22, 787–796. 10.1038/s41593-019-0364-9.

43. Li, N., Chen, T.-W., Guo, Z.V., Gerfen, C.R., and Svoboda, K. (2015). A motor cortex circuit for motor planning and movement. Nature 519, 51–56. 10.1038/nature14178.

44. Arroyo, B., Lemus, E.H., and Gutierrez, R. (2024). The flow of reward information through neuronal ensembles in the accumbens. Preprint at bioRxiv, 10.1101/2024.02.15.580379 https://doi.org/10.1101/2024.02.15.580379.

45. Musall, S., Kaufman, M.T., Juavinett, A.L., Gluf, S., and Churchland, A.K. (2019). Single-trial neural dynamics are dominated by richly varied movements. Nat Neurosci 22, 1677–1686. 10.1038/s41593-019-0502-4.

46. Steinmetz, N.A., Zatka-Haas, P., Carandini, M., and Harris, K.D. (2019). Distributed coding of choice, action and engagement across the mouse brain. Nature 576, 266–273. 10.1038/s41586-019-1787-x.

47. Juen, Z., Villavicencio, M., and Zuker, C.S. (2024). A neural substrate for short-term taste memories. Neuron 112, 277–287.e4. 10.1016/j.neuron.2023.10.009.

48. Eichenbaum, H. (2014). Time cells in the hippocampus: a new dimension for mapping memories. Nat Rev Neurosci 15, 732–744. 10.1038/nrn3827.

49. Pastalkova, E., Itskov, V., Amarasingham, A., and Buzsáki, G. (2008). Internally generated cell assembly sequences in the rat hippocampus. Science 321, 1322–1327. 10.1126/science.1159775.

50. Orhan, A.E., and Ma, W.J. (2019). A diverse range of factors affect the nature of neural representations underlying short-term memory. Nat Neurosci 22, 275–283. 10.1038/s41593-018-0314-y.

51. Williams, A.H., Poole, B., Maheswaranathan, N., Dhawale, A.K., Fisher, T., Wilson, C.D., Brann, D.H., Trautmann, E.M., Ryu, S., Shusterman, R., et al. (2020). Discovering Precise Temporal Patterns in Large-Scale Neural Recordings through Robust and Interpretable Time Warping. Neuron 105, 246–259.e8. 10.1016/j.neuron.2019.10.020.

52. Di Lorenzo, P., Chen, J.-Y., and Victor, J. (2009). Quality Time: Representation of a Multidimensional Sensory Domain through Temporal Coding. The Journal of neuroscience : the official journal of the Society for Neuroscience 29, 9227–9238. 10.1523/JNEUROSCI.5995-08.2009.

53. Coss, A., Suaste, E., and Gutierrez, R. (2022). Lateral NAc Shell D1 and D2 Neuronal Ensembles Concurrently Predict Licking Behavior and Categorize Sucrose Concentrations in a Context-dependent Manner. Neuroscience 493, 81–98. 10.1016/j.neuroscience.2022.04.022.

54. Talpir, I., and Livneh, Y. (2024). Stereotyped goal-directed manifold dynamics in the insular cortex. Cell Reports 43. 10.1016/j.celrep.2024.114027.

