## Supplementary figures and images for "Sequential and dynamic coding of water-sucrose categorization in rat gustatory cortices"

### Figure S1

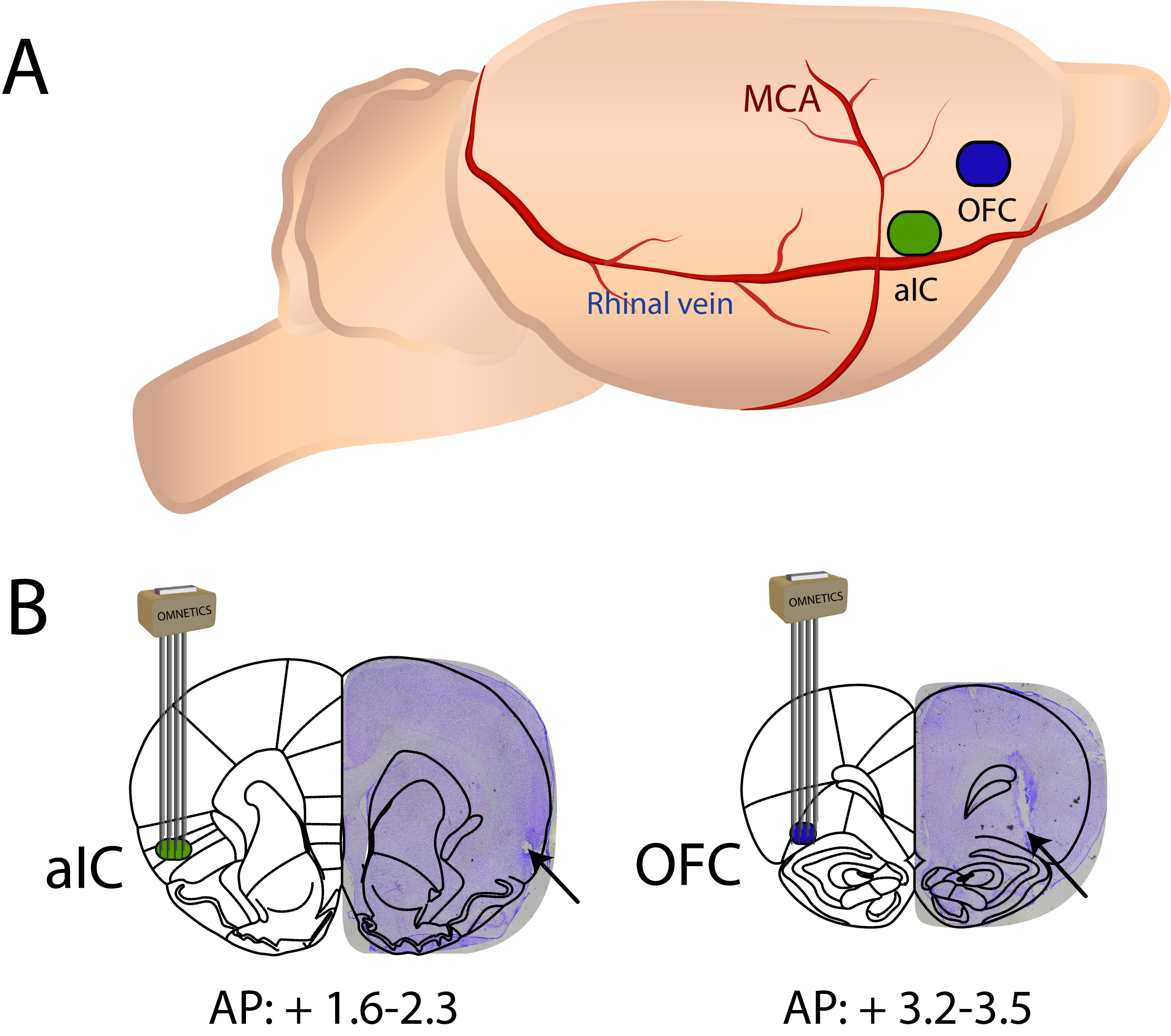

### Figure S2

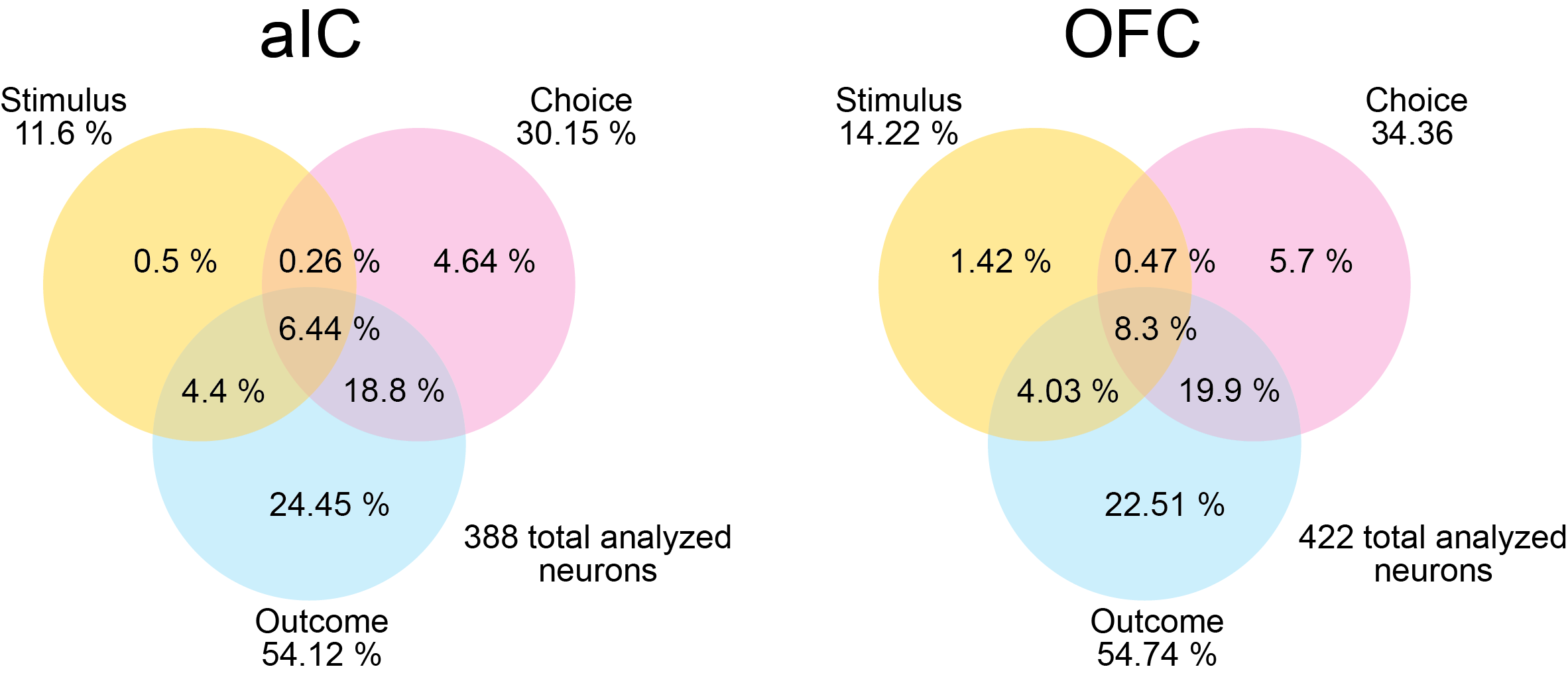

### Figure S3

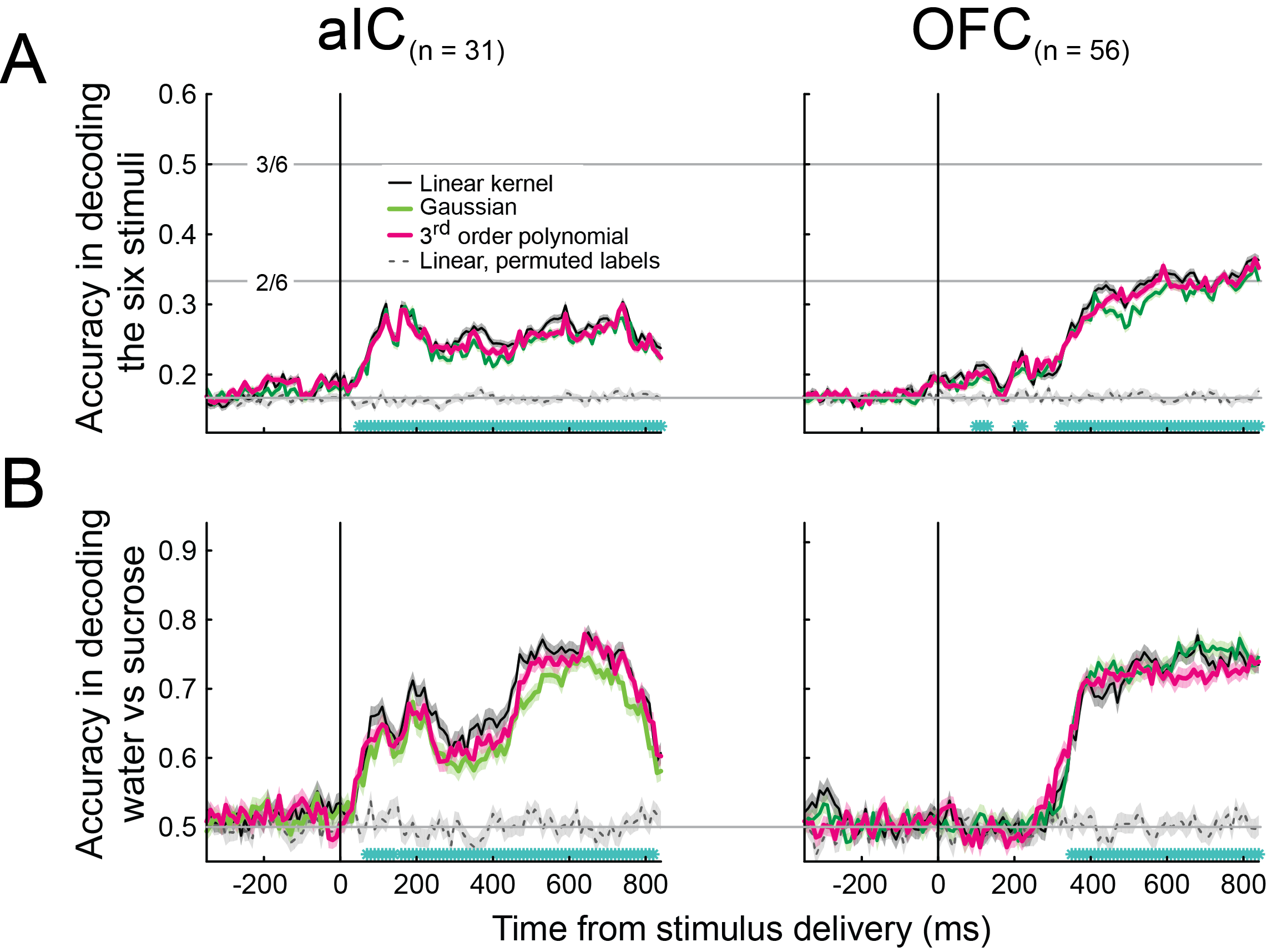

### Figure S4

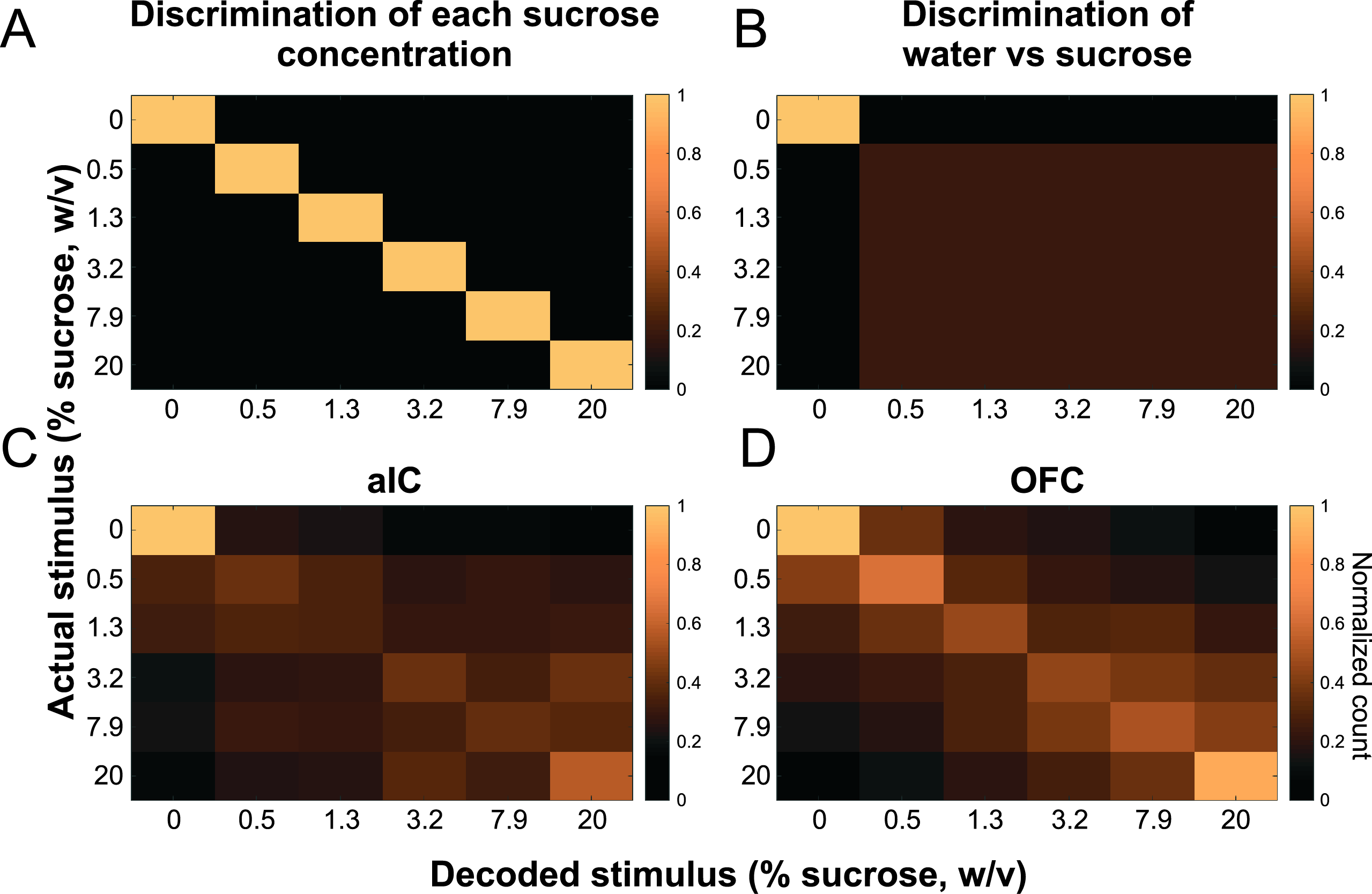

### Figure S5

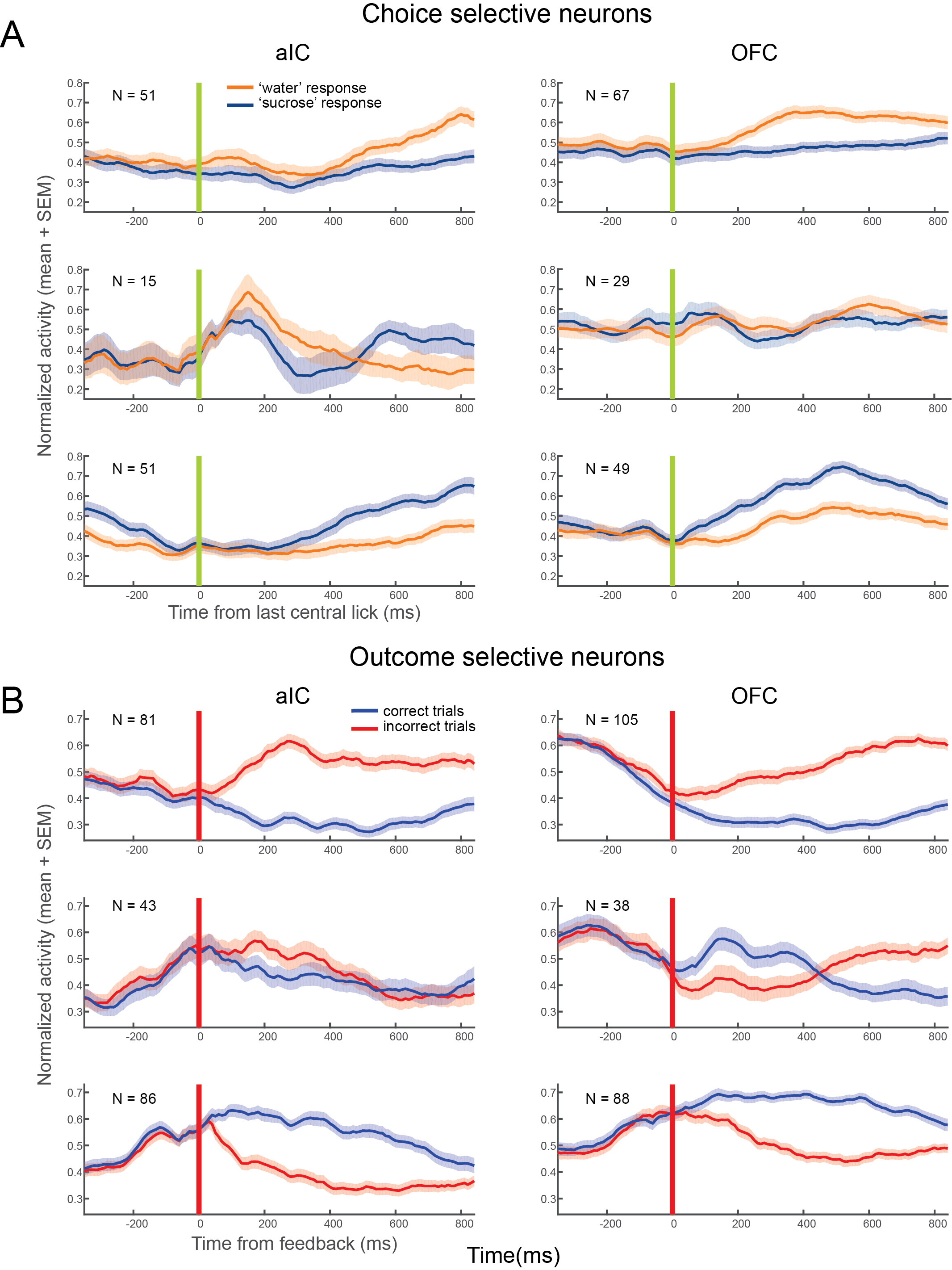

### Figure S6

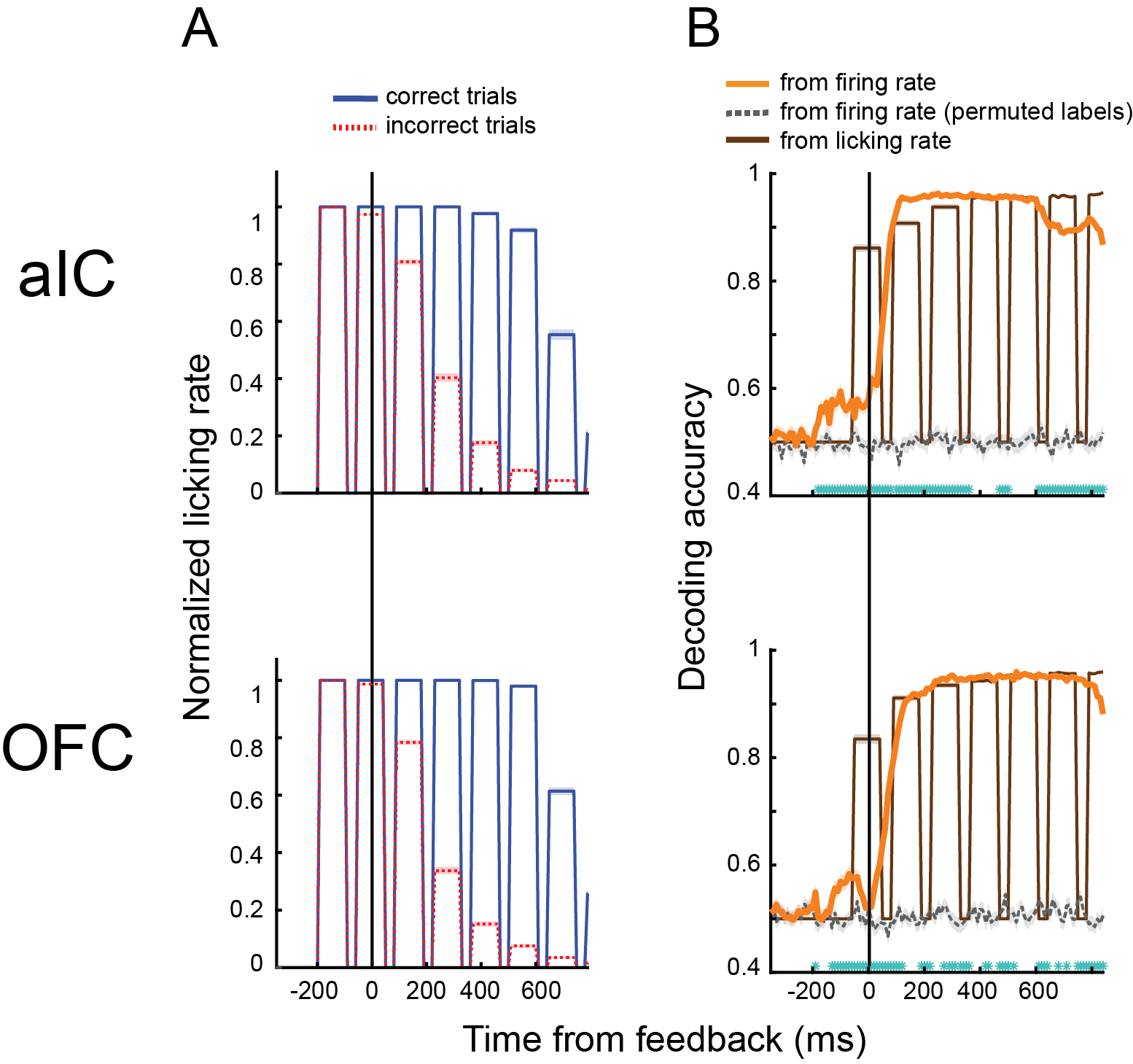
